# Quantifying transmembrane water exchange by diffusion NMR methods: From yeast cells to optic nerve ex vivo

**DOI:** 10.1101/2025.06.01.657299

**Authors:** Yuval Scher, Shlomi Reuveni, Yoram Cohen

## Abstract

Non-invasive measurement of exchange is paramount in different fields, ranging from material to biological sciences. Unaccounted exchange may even blur micro-structural or other characteristics of multi compartmental systems studied by MR methods. Despite the growing interest in diffusion-exchange studies of complex systems, comparative studies remain scarce. Most existing investigations have applied different diffusion MR methods to different biological samples under varying experimental conditions, making direct comparisons difficult. Moreover, the lack of a gold standard for exchange rate measurements further complicates efforts to validate and interpret results. To address these challenges, we employed two diffusion NMR-based methods—the constant-gradient pulsed field gradient (CG-PFG) and the recently introduced filter-exchange NMR spectroscopy (FEXSY)—to investigate apparent water exchange in yeast cells and optic nerves, both before and after fixation. We first evaluated the effect of the q-values on the extracted indices and then evaluated the repeatability and reproducibility of the measurements. The CG-PFG and FEXSY experiments were collected on the same sample to allow comparison of the results. The intracellular mean residence times (MRTs) (*τ*_*i*_) extracted from the log-linear fit of the CG-PFG NMR experiments were found to be 554 ± 6 ms and 337 ± 10 ms for yeast cells before and after fixation, respectively. The respective *τ*_*i*_ values extracted from the FEXSY experiments before and after fixation were found to be 368 ± 14 ms and 146 ± 24 ms, respectively. Despite the difference in absolute values of MRTs, the same qualitative behavior is observed in the two methodologies, and both could be analyzed using the bi-compartmental Kärger model. The same methodologies were then used to study exchange in the more complex porcine optic nerves. There, the bi-compartmental Kärger model analysis is shown to be inadequate. Extensive Monte Carlo simulations are used to narrow down on the most possible explanation, suggesting that optic nerves are multi-compartmental systems where not all spins are free to undergo exchange. Supporting theoretical calculations point to the existence of at least one additional non-exchanging restricted compartment. Thus, a tri-compartmental model is derived and used to analyze the data. The new model fits the data significantly better and results in dramatically different exchange rates when used on white matter (WM) data: CG-PFG experiments were found to be 730 ± 40 ms and 803 ± 16 ms for optic nerves before and after fixation, respectively. The respective *τ*_*i*_ values extracted from the FEXSY experiments before and after fixation were found to be 530 ± 125 ms and 387 ± 104 ms, respectively. These values are considerably lower than the values previously reported. Finally, we use simulations to show that the quantitative discrepancy between CG-PFG and FEXSY can be attributed, at least partially, to the difference in *T*_2_ values between the intra- and extracellular compartments. We thus encourage the pairing of exchange and spin-spin relaxation measurements in future studies. We end with a discussion on the current state of the diffusion-exchange, and in an attempt to put a spotlight on essential corner stones that are still missing despite the great advance of recent years: experimental standardization, method comparison, and adequate modeling.

## 1. INTRODUCTION

Diffusion magnetic resonance (MR) spectroscopy and imaging (d-MRS and d-MRI, respectively) have been used extensively to study, non-invasively, the microstructural characteristics of porous materials and different biological tissues in vivo, with special emphasis on neuronal tissues and the central nervous system (CNS) [1–4]. Understanding microstructural features is crucial, as they often play a key role in determining functional properties across various systems. For example, it is known that the size of the pores affects the catalytic activity and storage capacity of many porous materials [4] and the size of the axon affects, at least in myelinated nerves, the speed of neuronal transmission [5]. Abnormal axon sizes were also found to be associated with different neurological disorders such as amyotrophic lateral sclerosis (ALS) and multiple sclerosis (MS) [6, 7].

In the last two decades, much effort has been devoted to extract microstructural information from tissues and organs non-invasively [3, 8–18]. However, in multi-compartmental systems, exchange across boundaries and membranes may blur the differences in the measured NMR characteristics of the different compartments. As a result, most microstructural studies have assumed impermeable membranes, effectively neglecting exchange altogether [3, 8–17]. At the same time, water exchange and membrane permeability are crucial parameters in various contexts, from understanding membrane-based processes to detecting and monitoring disease. Notably, changes in cell membrane permeability are known to be biomarkers for certain pathological conditions [19– 23]. This underscores the need for new, robust, and flexible non-invasive methods to accurately quantify water exchange across membranes [24].

One of the earliest attempts to characterize exchange with pulsed field gradient (PFG) diffusion NMR experiments was proposed by Kärger [26–29]. This model, which is based on the extension of the bi-compartmental Stejskal-Tanner equation [30, 31], has been used to study exchange in different chemical systems and in different cells [27, 28, 32–34]. It assumes that the two exchanging populations exhibit Gaussian (free) diffusion; an assumption which generally does not hold in cell systems and many other biological tissues. Price and coworkers modified the Kärger model to account for systems in which one of the diffusing components exhibits restricted diffusion in spheres [35]. The Leibfritz group used the so-called constant-gradient pulsed gradient stimulated echo (CG-PGSTE [31], see Figure 1(a)) approach to study diffusion exchange in neuronal cells and tissues [36–38]. Meier et al. modified the Kärger model to account for systems in which one of the diffusing components is restricted in cylinders [37, 38]. Other more specialized models for the analysis of PFG experiments were also considered [39, 40]. For example, a comprehensive diffusion MRS study of bovine optic nerve was conducted by Stanisz et al. [40]. They used a tri-compartmental model: axons (prolate ellipsoid), glial cells (spheres), and extracellular medium.

**FIG. 1.**
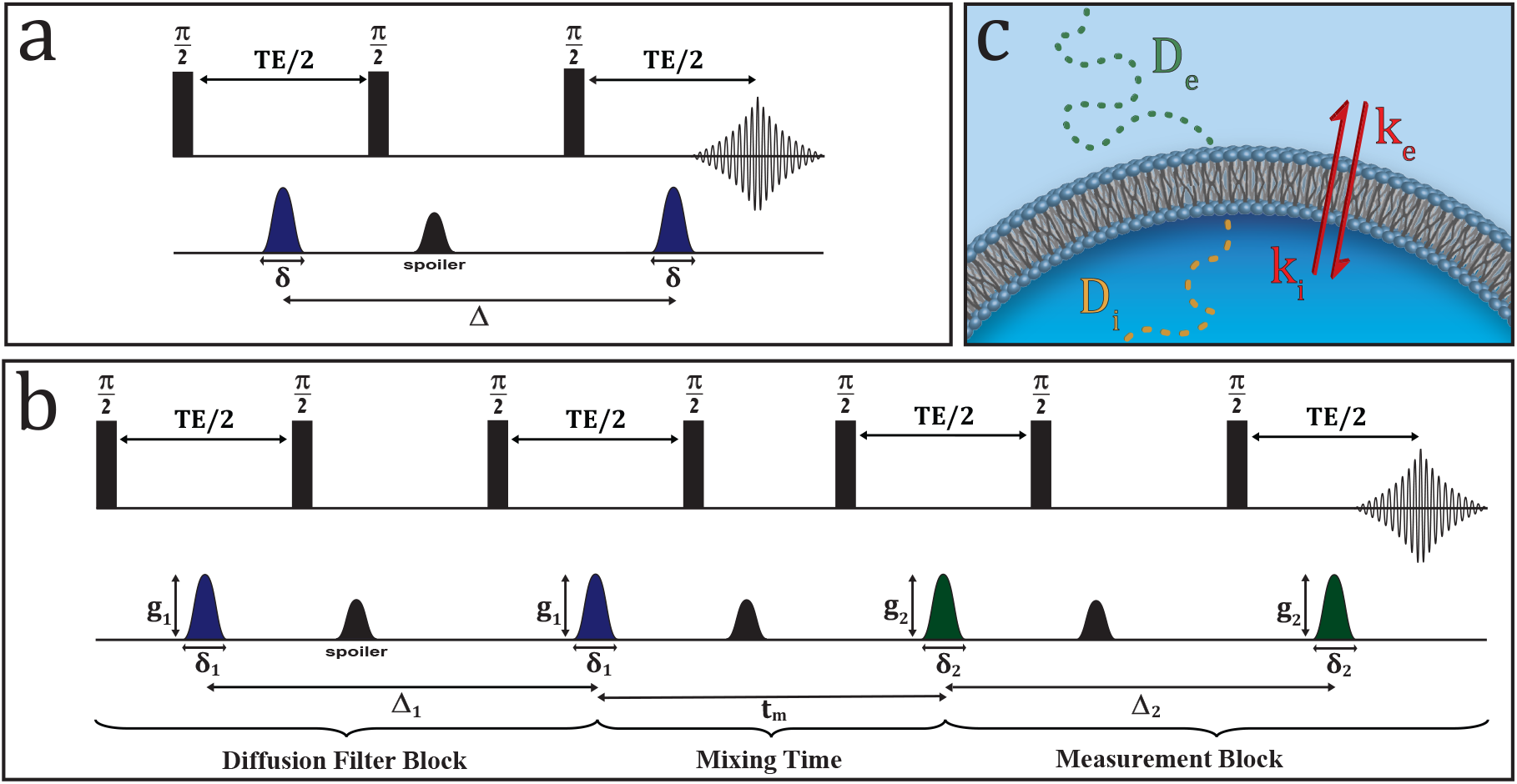
(a) The PGSTE NMR sequence [31], (b) the FEXSY sequence [46], and (c) a schematic representation of a bi-compartmental cell system, reproduced from Ref. [48].

In parallel to the introduction of the CG-PGSTE experiments for exchange measurement, Callaghan and coworkers developed the two-dimensional diffusion exchange NMR spectroscopy method named 2D-DEXSY [41–44] for measuring exchange. In this approach, the pulse sequence is composed of two PFG NMR blocks separated by a mixing time (*t*_*m*_). The pulse gradient pairs are increased independently to form a 2D matrix. In the DEXSY sequence, each block records the displacement at a different given time. Therefore, molecules exchanging diffusion sites during the time interval between the two diffusion blocks appear as cross peaks on a computed 2D correlation map. One of the main advantages of this approach is that producing the correlation map does not require any model, thus allowing for the observation of exchange between multiple sites relatively easily. However, for obtaining quantitative measures, one must repeat the 2D experiments at different mixing times, which renders this approach extremely demanding in terms of scan time. Recently, Benjamini et al. suggested a sampling and processing strategy that significantly reduces the time required to perform such experiments [45].

Åslund et al. proposed to fix the gradient intensity of the first diffusion block of a 2D-DEXSY experiment, thus transforming it into a constant preparation module which in turn reduces the dimensionality of the DEXSY experiment (see Figure 1(b)) [46]. In this variant, a preparation module serves as a displacement-based filter block that precedes a conventional PFG NMR experiment. The filter block selectively attenuates the signal of the fast diffusing populations in the explored multi-compartmental or multi-component systems. Å slund et al. showed that by changing the *t*_*m*_ between the filter block and the measurement block in repeated experiments (where the filter block is kept constant), it is possible to measure the exchange between the slowly diffusing population and the filtered population. The analysis of such experiment, called later diffusion filter-exchange NMR spectroscopy (FEXSY), was extended, with the aid of 2D diffusion-relaxation measurements, to two-compartmental systems in which the diffusing spins in the different compartments also differ in their relaxation rates [47]. Recently, Scher et al. demonstrated that it is possible to obtain the exchange rate from a partial FEXSY data set which was called Constant Gradient FEXSY, or CG-FEXSY [48].

The FEXSY methodology was also extended to MRI and was termed filter-exchange MR imaging or FEXI [49–51]. There, using a phenomenological equation, apparent exchange rates (AXRs) have been measured and even imaged in different systems using both pre-clinical and clinical MRI scanners [49–52]. FEXI was elegantly used to study indirectly over-expression of the urea transporter both in cells and in vivo [53]. Recently, a modified protocol termed TEXI was introduced for thin imaging slices, which are more commonly used in preclinical settings [54].

Finally, there is a continuous endeavor on part of the Basser lab to reduce the DEXSY experiment time and to extract more structural and dynamical information from it, while also making a case for the use of static-field gradients in lieu of pulsed-field gradients [45, 55–60]. We have already mentioned by passing the processing strategy utilized by Benjamini et al. [45]. A later approach by Cai et al. used a reparameterizaion that utilizes the symmetry of the 2D-correlation map, originally proposed by Song et al. for analysis of 2D relaxation-MR experiments [61], to further reduce the required data points to be collected [56]. In fact, their approach leads to a novel way of extracting the exchange rate from the 2D-DEXSY correlation map: After reparameterization, they show that only four points per *t*_*m*_ are required to calculate the exchanging fraction. Then, the dependence of the exchange fraction on *t*_*m*_ can be fitted to obtain the exchange rate [57]. While this method is still more time-consuming than CG-FEXSY sampling, which requires only a single point per *t*_*m*_ (divided by a normalization data point) [48], it is expected to work better in tissues where there is further weighting by the spin-lattice relaxation of a non-exchanging population, if such exist (and in fact, it offers a way to measure its relaxation rate) [57]. Note, however, that this method does not account for a difference in the relaxations of the exchanging pools, as was done in the aforementioned procedure by Eriksson et al. to account for the spin-spin relaxation [47].

Last year, Diffusion Exchange Ratio (DEXR) analysis was introduced [60]. It seems to use the same procedure as CG-FEXSY, albeit with static-field gradients instead of pulsed-field gradients. The authors acknowledge the similarity, but state that the FEXSY model is fundamentally different in that it looks at an apparent diffusion coefficient (ADC) recovery [60]. Indeed, it is true that a simple phenomenological fit of ADC recovery was introduced as well [49], but the original works of FEXSY and CG-FEXSY dealt directly with the signal attenuation [46, 48]. Hence, consideration of models that go beyond exchanging free-diffusing population is possi-ble (as was done for CG-PFG [35–38]), and will be discussed herein. In light of these very new findings, it is clear that a direct comparison study between the plethora of DEXSY-derived methods is in place. A first step in this direction was recently taken by Ordinola et al. [62].

The state of affairs is such that a non-expert researcher aiming to measure exchange rates with d-MRS methods will face an overwhelming number of acronyms and abbreviations, and no organized review to aid. A partial list would be: CG [37], DEXSY [43], FEXSY [46], CG-FEXSY [48], FEXI [49], TEXI [54], NEXI [63], REEDS-DE [58], DEXR [60], SANDIX [64], INDIANA [65], LLPS REDIFINE [67], EXCHANGE [22], DKI(*t*) [68] and JOINT [69]. Indeed, every model or acquisition scheme has its own nuanced difference, and some models are specialized to explore a specific type of tissue or process. Yet the risk of redundancy rapidly increases. In fact, we can roughly separate the methods into two diffusion-based approaches: single vs. double diffusion encoding d-MR methods [70]. In each, the extent to which the experimental parameter space is sampled varies (as higher parameterization of the tissue model often requires more data collection). Similarly, all works use either the Kärger model or the proper modified version of it. In fact, theoretically one can try to do without the well-mixing assumption of the Kärger model and solve for the overall signal attenuation directly from the diffusion propagator in the fashion of Grebenkov et al. [71]. However, such a theory was neither experimentally tested, nor was the (plausible) validity of the well-mixing assumption.

Despite the surge in studies devoted to diffusion-exchange in complex systems, i.e., systems in which at least one of the exchanging components exhibits non-Gaussian diffusion, there are only very few comparative studies where the same system was simultaneously studied by two different diffusion-based approaches [67, 68, 72]. In most of the diffusion exchange studies reported to date, different cells and tissues were investigated using different diffusion MR methods and approximations. This was generally done with different diffusion weighting and at different experimental conditions (temperature, preparation, and concentrations) thus precluding direct comparison. This problem is further exacerbated by the lack of a gold standard for exchange rates. Clearly, the situation here is even more problematic than when attempting to obtain detailed microstructural information from MR data. There one can use phantoms or simple biological systems whose size and shape are known a priori. In more complex systems one may still rely, at least partially, on histology and electron microscopy [15–17, 73–82] to challenge the MR results. Yet, similar benchmarks for exchange rates are currently lacking.

In the present study, we directly compare two of the most prevalent diffusion NMR-based approaches used to study exchange rates in biological systems, namely the CG-PGSTE and FEXSY methods. Exchange rates in yeast cells and optic nerves ex vivo were studied both before and after fixation. CG-PGSTE and FEXSY NMR experiments were performed on the same sample consecutively to allow comparison of the results, and in some cases the samples were followed for a period of up to 12-16 hours. This was done to show the reproducibility and repeatability of the extracted indices, both in terms of the measurements and the interrogated samples.

The methods agree qualitatively when comparing different systems. However, the two methods do not agree quantitatively, the FEXSY exchange rate is ∼1.5 − 2.5 higher than the CG-PGSTE. Markedly, we show that fixation leads to considerably faster exchange, which corresponds to greater membrane permeability. This is well captured by both methods: In yeast cells, fixation leads to an increase in permeability by a factor of ∼2.5 according to FEXSY and of ∼1.5 according to CG-PGSTE (suggesting that FEXSY is more sensitive). In optic nerves, however, the effect is insignificant. As mentioned, exchange acts to blur the differences between different compartments, and so a higher exchange rate can lead to a larger error in the determination of, for example, the compartment size or the relative fractions of spins in the two domains (that is, if exchange is not included in the model) [69, 83]. As tissue fixation is a common practice, this result suggests that caution is needed when reporting the compartment size or fractional populations of the intra- and extracellular compartments of fixed tissues. We demonstrate this effect using a set of Monte Carlo simulations.

We chose to study systems of increasing complexity. We first studied yeast cells and then used the same methodologies to study optic nerves, which can serve as a model for the white matter (WM). Evaluating the validity of the bi-compartmental model, we show that it is adequate when studying yeast cells. Unfortunately, we show that the bi-compartmental model breaks down when considering the optic nerve, despite being used routinely in the analysis of WM.

We further challenged the experimental findings with Monte Carlo simulations of a sample akin to yeast cells. There are many factors that are present in vivo or ex vivo experiments but are unaccounted for in existing models, e.g., the spin-spin relaxation effects discussed above. The series of idealized in silico experiments that we performed provides a look at the inherent difference between the two methods without these effects, allowing us to isolate the effects of diffusion exchange. In silico, we find that both FEXSY and CG-PGSTE work very well in extracting the exchange rate, which narrows down the possible set of explanations for the experimental discrepancy. We simulated a range of cell densities, with the upper limit being hexagonal close-packed (HCP) lattice. In the HCP case, the extracellular fraction is actually more restricted than the intracellular fraction, similar to the case of the optic nerve. Surprisingly, FEXSY described the simulated data from the HCP lattice well. We thus conclude that, while the density can affect the accuracy of the FEXSY measurement (especially when the extracellular compartment becomes restricted itself), this by itself cannot explain the failure of FEXSY in describing the optic nerve. It is more likely that the bi-compartmental assumption is invalid. For example, there might be a third population of spins not partaking in exchange. While the effect of a third compartment is immediately apparent in the FEXSY experiment, the effect is more subtle in the CG experiment. For a truly bi-compartmental system, changing the constant gradient used for measurement does not effect the result. This however is no longer the case if we add the third population.

To further test the hypothesis of a third non-exchanging population, we derived tri-compartmental models for CG-PGSTE and FEXSY. The tri-compartmental model can explain the breakdown in the analysis, where the current analysis methods being employed in white matter are likely to lead to erroneous values being reported. In turn, these theoretical calculations revealed, through the mismatch between the CG-PGSTE WM data and the bi-compartmental model, that the required additional population must be restricted. We show that for the analysis of both CG-PGSTE and FEXSY experiments correction of the model is simple and straightforward, and that the exchange information can be achieved by a novel fit of the data to attain *k*_*i*_.

Lastly, we provide a comprehensive discussion of the results, how do they compare to existing literature, and their implications.

The navigation map for this paper is as follows. In Sec. 2 A we derive and discuss the bi-compartmental models, and in the following subsections 2 B-2 F we report the samples preparation procedures, the hardware used, and the experimental methods. Finally, in Sec. 2 G we explain the algorithm for the Monte Carlo simulations employed in this paper. In Sec. 3 we present and discuss the results, using the bi-compartmental analysis. In Sec. 4 we derive the tri-compartmental model, perform the corresponding simulations, and re-analyze the data from the optic nerves. In Sec. 5 we show that a difference in intra- and extracellular *T*_2_ values can explain, at least partially, the quantitative discrepancy between the two meth-ods. In Sec. 6 we discuss all of the findings reported in this paper and how they relate to previous literature. We further survey some interesting recent developments. Importantly, in Sec. 6 E we discuss WM modeling. Finally, we conclude in Sec. 7.

## 2. MATERIALS AND METHODS

### A. Modeling

The PGSTE sequence in Figure 1(a) is a conventional d-MRS technique [31], where the location of the diffusing spins is tagged on two instances separated by a diffusion period Δ. The tagging is performed by applying pulsed-field gradients of duration *δ*, which exert a spatially dependent phase. The phase acquired by each spin during the first gradient pulse is inverted by application of RF pulses during the diffusion time, such that it cancels with the phase acquired during the second identical gradient pulse only if no translation occurred during that time. More explicitly, under the short gradient pulse (SGP) approximation, i.e., *δ* → 0 and Δ ≫ *δ*, the effect of a pulsed-field gradient (*g*), on the phase of a single spin at position **r** is given by [2]

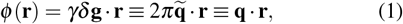

where *γ* is its gyromagnetic ratio and q is given in units of *rad* · *m*^−1^. Considering a spin at position **r**_0_ during the first gradient pulse and at position **r** during the second gradient, the overall accumulated phase in such a PFG experiment is then given by

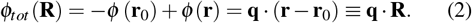

Under these conditions, the attenuation of the echo intensity due to diffusion can be written as an ensemble average of the phases [1, 2]

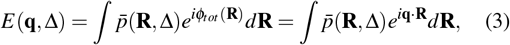

where the phases are weighted by the averaged diffusion propagator

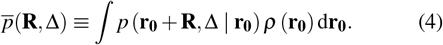

Here, the diffusion propagator *p* (**r**_**0**_ + **R**, Δ | **r**_0_) represents the probability that a particle with initial position **r**_0_ will undergo a displacement **R** during the diffusion time Δ. The average diffusion propagator is then given by integrating the diffusion propagator over the initial density *ρ*(**r**_0_) as presented in Eq. (4), which in fact describes the displacement distribution of the spins during the diffusion time [1, 2].

In the case of freely diffusing spins, the position at time Δ is independent of the initial position, hence *p*(**R**, Δ) reduces to 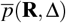, which is the well-known Gaussian distribution 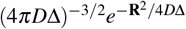, where *D* is the diffusion coef-ficient. Indeed, inserting the Gaussian diffusion propagator of free-diffusing spins into Eq. (3) yields the well-known Stejskal-Tanner equation (N.B., freely diffusing spins in the limit *δ* → 0 and Δ ≫ *δ*)

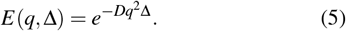

Equation (5) implies that the diffusion coefficient can be extracted from a log-linear fit. Varying *q* while keeping all time intervals constant yields the constant time (CT) experiment. Varying Δ while keeping the q-value constant yields the constant gradient (CG) experiment [36–38].

For bi-component systems, one can extend Eq. (5)

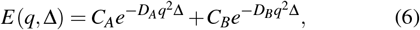

where *C*_*A*_ and *C*_*B*_ represent the relative population of spins having diffusion coefficients *D*_*A*_ and *D*_*B*_, respectively. Note that the above extension assumes Gaussian diffusion, neglects the effect of relaxation, and, more importantly in the context of this study, assumes no exchange between the two populations. It is well-known, however, that for a particle trapped in a compartment, the log-linear fit described above results in an ADC which is smaller than the actual intrinsic free-diffusion coefficient *D*.

It is convenient to define the dimensionless variable *ξ* = 2*D*Δ*/d*^2^, i.e., the ratio between the square of the displacement during the diffusion time and the square of the characteristic size of the compartment [1, 2]. Note that *ξ* is larger than 1 when diffusion is restricted. For very large *ξ* ≫ 1, the root mean square (rms) displacement reaches its upper restricting value, which corresponds to the characteristic size of the compartment. In these cases, the diffusion propagator is merely the equilibrium density as the spins sample the entire compartment during the diffusion time and hence we can rewrite Eq. (4) as

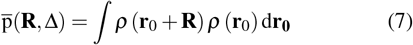

Equation (7) is a density autocorrelation function, which implies that the decay of the echo intensity for restricted diffusion in various simple geometries can be calculated and predicted based on the knowledge of *ρ*(**r**_0_). This was demonstrated by Callaghan and others [1, 2, 84].

As a pertinent example, consider diffusion restricted to a spherical compartment with diameter *a*, long diffusion time such that *ξ* ≫ 1 and the application of a short gradient pulse in the x-direction. In this limit we have [85]

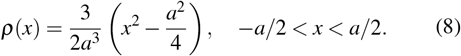

Plugging into Eq. (3) we obtain

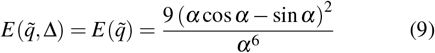

Where 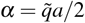. Note that the plot of 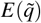 as a function of *α* exhibits a diffraction pattern with a node at *α* ≈ 3*/*4 [85].

In the yeast cell sample, as presented in Figure 1(c), the diffusion signal attenuation can be described by the bicompartmental model (Eq. (6)) using the notation presented in the following equation

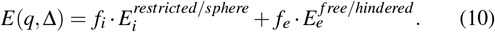

In this equation, 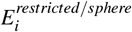 and 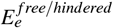 represent intracellular and extracellular components having an intra- and extracellular fraction (*f*_*i*_ and *f*_*e*_), respectively. For convenience, the superscripts will be omitted henceforward. Note that such a model assumes monodispersity in size, no orientation dispersion, equal relaxation rates for all populations and no coupling or exchange between the two populations.

As stated, the Kärger model was the first attempt to introduce exchange into the bi-compartmental model assuming, however, Gaussian diffusion for the two diffusing components or compartments and well-mixing. In the Kärger model, the exchange terms are added as a coupling between the populations. Using the matrix notation, for two populations exhibiting Gaussian diffusion we can write [28, 34]

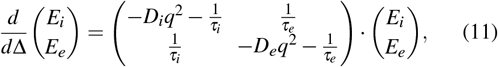

where *τ*_*i*_ and *τ*_*e*_ stand for the mean residence time (MRT) in the intra- and extracellular compartments, i.e., the reciprocal of the exchange rates *k*_*i*_ and *k*_*e*_, respectively. Similarly, *D*_*i*_ and *D*_*e*_ are the intrinsic intra- and extracellular diffusion coefficients.

We assume that initially

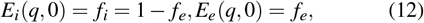

such that equilibrium is maintained (influx to the cell is equal to the efflux)

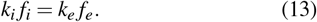

The matrix can then be transformed to an uncoupled coordinate system and solved to give a bi-exponential expression with the form of Eq. (6), but with effective diffusion coefficients (*D*_*A*_ and *D*_*B*_) and effective fractional populations (*C*_*A*_ and *C*_*B*_).

For our purposes, it is enough to examine the expression for *D*_*A*_ and *D*_*B*_

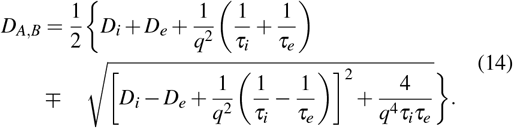

In a similar manner, Price et al. solved a modified Kärger model where one of the populations is restricted to a sphere (*ξ >* 1, under the SGP approximation) [35]. While Eq. (14) retains its form (up to setting *D*_*i*_ = 0, which is intuitive), the effective fractional populations are different. Meier et al. adapted Price’s derivation to fit cylindrical compartments, using the Gaussian phase distribution approximation [37, 38].

Extracting all five parameters from the three free apparent parameters of the bi-exponential fit is relatively demanding. Instead, it is customary to consider the slow exchange limit, that is, *q*^2^*D*_*e*_ ≫ *k*_*i*_, *k*_*e*_, in which *D*_*A*_ can be approximated as

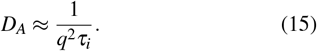

Meier et al. introduced an even simpler method [37]. If the difference between the ADCs of the two populations is significant, i.e., the faster ADC_*e*_ is much larger than the slower ADC_*i*_, the signal from the fast-diffusing population will attenuate much faster. Thus, we can find sufficiently long diffusion times, in which the faster diffusing fraction is completely suppressed, while some signal from the slow fraction is retained. In fact, the slow exchange limit means that the fast diffusing population must attenuate faster than the exchange rates, i.e., *q*^2^*D*_*e*_ ≫ 1*/τ*_*i*_, 1*/τ*_*e*_. Here clearly high q-values lead to a more effective selective attenuation of the fast diffusing population. If one of the populations is restricted, it is even easier to create such differences between the ADCs of the two compartments by increasing the diffusing time. Under the slow-exchange limit and a suppressed fast diffusing population, the slope of a CG diffusion NMR experiment and the MRT in the restricted compartment (*τ*_*i*_) are related by

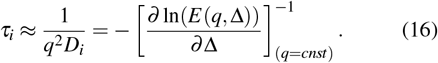

This elegant method was used by the Leibfritz group to estimate the MRTs in neuronal cells and CNS tissues [36–38].

So far for the CG-PFG experiment. When the FEXSY experiment (Figure 1(b)) [46] is applied to the system shown in Figure 1(c) with a zero mixing time *t*_*m*_, it is clear that the signal of the fast diffusing population, *f*_*e*_(0), will be suppressed to some extent. The signal of the slow diffusing population, *f*_*i*_(0), will also be suppressed, but to a much lesser extent. Now, increasing the *t*_*m*_ will allow the fractional populations *f*_*e*_(*t*_*m*_) and *f*_*i*_(*t*_*m*_) to change, since increasing the mixing time enables more water molecules to diffuse across the membrane and change their affiliation. At very long mixing times, *f*_*e*_(*t*_*m*_) and *f* (*t*) should approach their equilibrium values, 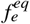 and 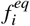, respectively. In the FEXSY experiment, assuming well-mixing, the fractional population at a given *t*_*m*_ follows

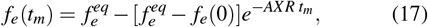

where 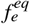 is the fractional population in equilibrium and AXR is the Apparent eXchange Rate. If the investigated system is a pure bi-compartmental system, AXR and *k* = *k*_*e*_ + *k*_*i*_ should be the same [46, 48, 50, 72] and so in this work we will use AXR and *k* interchangeably. For a system in equilibrium (i.e., *k*_*i*_ *f*_*i*_ = *k*_*e*_ *f*_*e*_, as in Eq. (13)) we obtain

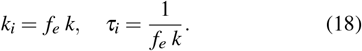

From the relations in Eqs. (16) and (18) it follows that *τ*_*i*_ can be calculated so that the results of the CG and FEXSY experiments can be compared. Note also that 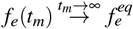 and therefore, theoretically, the attenuation of a PGSTE experiment or a FEXSY experiment with *g* = 0 in the filter block is equivalent to a FEXSY experiment in the limit of *t*_*m*_ → ∞, neglecting relaxation. There, the echo intensity attenuation is simply the superposition of two attenuations, weighed by their fractional populations

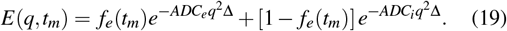

where the effect of restriction is incorporated into *ADC*_*i*_ [86]. Considering an intracellular spherical compartment, a modified version of Eq. (19) with the intracellular signal estimated using the SGP approximation (see Eq. [3] in Ref. [87]) does not result in the measurement of a different exchange rate or fractional populations, but in principle provides an easy way to further extract the radius of the cells by replacing the second exponent (19) with the proper infinite sum. If *ξ* ≫ 1, we retain only its first term, which is given by Eq. (9), and Eq. (19) is modified to

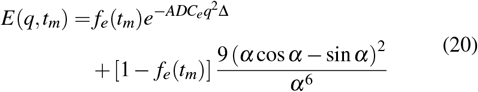

where 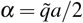. However, note that in FEXSY we often keep Δ relatively small to minimize exchange during the measurement block, so that, to a good approximation, exchange takes place mostly during the mixing time. Therefore, the limit *ξ* ≫ 1 is usually not reached and higher orders are needed for an accurate estimation of the radius. Essentially, there is a trade-off between the accuracy of measuring the exchange rate and the radius.

Lasič, Nilsson et al. introduced a simpler phenomenological fit of the FEXSY data [49]. In this approach, an ADC is extracted from the low q-limit of each of the FEXSY experiments. The ADC in an experiment without a preceding filter is simply

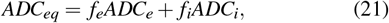

and the ADCs of the filtered experiments are given by

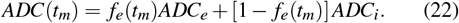

An analogous expression to Eq. (17) can then be defined in terms of apparent ADCs

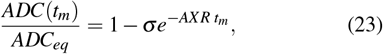

where *σ* = 1 − *ADC*(0)*/ADC*_*eq*_ is the filter efficiency and AXR is the apparent exchange rate. Therein, the inverse of AXR is termed apparent MRT.

### B. Yeast cells preparation

A colony of the BY4741 strain of Saccharomyces Cerevisiae was grown on a petri dish containing agar kept at 4°C. The NMR samples were prepared by taking a small amount of yeast cells from the colony and suspending it in 45 mL of Synthetic Defined Medium. The cell suspension was then left overnight in an incubator shaker (30°C, 140 RPM). During the night, an amount suitable for the preparation of a 5 mm NMR sample had grown. The yeast cells were harvested from the suspension by centrifugation (as in Ref. [88]) and removal of the supernatant and then washed (re-suspended and harvested) three times with double-distilled water. Finally, the yeast cells were transferred to a 5 mm NMR tube. The NMR tube was centrifuged, and the supernatant was removed. The samples of the fixed yeast cells were prepared in the same manner, with an additional fixation step preceding the transfer to the NMR tube. The fixation was carried out by immersing the yeast cells in 4% formaldehyde PBS solution for 15 min. The yeast cells were then washed twice with PBS solution and transferred to the NMR tube.

### C. Optic nerves preparation

Five-month-old porcine optic nerves were dissected at the Institute of Animal Research Lahav (Kibbutz Lahav, Israel). The optic nerves were immediately immersed in a saline or formalin solution and immediately sent to TAU. NMR experiments of the non-fixed nerves started about 4 hours after nerve incision. The nerves that were taken out from the saline solution were gently dried, then carefully mounted into a 4 or 5 mm NMR sleeve and transferred into an 8 mm NMR tube filled with Fluorinert (Sigma, Saint-Louis, MO, USA). For the fixed samples, nerves were kept in formalin solution for 24 hours at room temperature and in PBS solution for an additional 24 hours. The nerves were then gently dried and carefully mounted into the 4-5 mm NMR sleeve which was then transferred into a Fluorinert filled 8 mm NMR tube.

### D. Hardware

All NMR experiments were performed on a Bruker 9.4 T WB NMR spectrometer equipped with a Bruker Micro5 probe capable of producing pulse gradients of up to 300 G/cm with 60 mA amplifiers. The probe temperature was kept constant at 25°C. Sine-shaped pulsed-gradients were used in all the performed diffusion NMR experiments and the duration of the *π/*2 RF pulses was about 15 *µ*s. The NMR sequences used in this study are depicted in Figure 1.

### E. Yeast Cells MR experimental parameters

#### i. Constant gradient PGSTE (CG-PGSTE)

The CG-PGSTE NMR experiments (Figure 1(a)) were performed with a TR/TE of 2000/8 ms and 4 dummy scans. The sine-shaped gradient pulse duration (*δ*) was 3.14 ms (i.e., *δ*_*e f f*_ of 2 ms) and the diffusion time (Δ) was incremented from 8 to 800 ms in 30 steps. The PGSTE NMR experiment was repeated with different constant gradient strengths (*g*) of 15, 30, 50, 100, 150, and 250 G/cm. A spoiler gradient was used during the z-storage period with *δ*_eff_ of 2 ms and *g* of 150 G/cm. The signal-to-noise ratio (SNR) for a typical CG-PGSTE NMR experiment on yeast cell samples with *g* of 100 G/cm and Δ of 8 ms was ∼5000. Note that for the logfit analysis of the CG-PGSTE data, the highest 10 data points (of the 30 acquired) were used.

#### ii. Filter exchange spectroscopy (FEXSY)

The FEXSY experiments (Figure 1(b)) were performed with a TR/TE of 2000/8 ms and Δ*/δ* of 13/3.14 ms in both diffusion blocks. Four dummy scans were used. In the filter block CG of 150 G/cm were used. For the measurement block, the gradient strength was incremented from 5 to 150 G/cm in 30 equally spaced steps. Mixing times (*t*_*m*_) of 8, 25, 30, 50, 100, 150, 200, 250, 300, 400, 500, 750, 1000 and 1250 ms were used. The no filter experiment was a PGSTE NMR experiment with the same parameters as the measurement block. Spoiler gradients were used during the z-storage periods with *δ*_eff_ of 1.27 ms and *g* of 20 G/cm. The SNR for a typical no-filter FEXSY or PGSTE experiments on yeast cell samples, with a g-value of 5 G/cm for the measurement block was ∼25000. For the quantitative analysis, only the data with mixing times of up to 750 and 400 ms were used for the fresh and fixed yeast cell samples, respectively. This was done due to the low SNRs of the data with higher *t*_*m*_s in each sample.

### F. Optic nerves MR experimental parameters

#### i. Constant gradient PGSTE (CG-PGSTE)

The CG-PGSTE NMR experiments were performed with a TR/TE/*δ* of 3000/8/3.14 ms and four dummy scans. The Δ was incremented from 11 to 1500 ms in 75 steps and the experiments were repeated with g-values of 15, 30, 50, 100, 150, and 250 G/cm. The SNR for a typical PGSTE experiment on nerve samples with g of 100 G/cm and Δ of 8 ms was 250.

#### ii. Filter exchange spectroscopy (FEXSY)

The FEXSY experiments were performed with a TR/TE/Δ/*δ* of 3000/8/35/3.14 ms in both diffusion blocks (i.e., the filter and measurement blocks) and four dummy scans. In the filter block a constant gradient of 285 G/cm were used. For the measurement block the gradient strength was incremented to 250 G/cm and in cases to 285 G/cm in 32 or 75 equally spaced steps, respectively. Mixing times of 8, 25, 30, 50, 100, 150, 200, 250, 300, 400, 500, 750, 1000 and 1250 ms were used. Note that in cases where gradients were incremented to 250 G/cm we could only include data for *t*_*m*_ of up to 750 ms because of the poor data quality at higher mixing times under these experimental conditions. The no-filter experiment was, again, a PGSTE NMR experiment with the same parameters as the measurement block. SNR of ∼4000 was found for a typical non-filter FEXSY experiment of nerve sample collected with g-values of 5 G/cm for the measurement block.

### G. Simulations

To further validate the experimental methods, Monte-Carlo simulations were performed using MATLAB R2021b (The MathWorks, Inc.). The code and files required to reproduce all the analysis in this manuscript are available in the Github repository, and as supplemental material.

Inside a domain (a rectangular cuboid), the code generates homogeneous spheres of a desired packing and size: (1) Random packing; (2) Layers of spheres, where the centers of the spheres lie on lines parallel to the axes; and (3) Hexagonal close-packed (HCP) lattice. The fractional volume of the spheres in the containing space is calculated.

The second step is to distribute the initial positions of the spins inside the containing space. Here, the spins are randomly distributed in the containing space (the code allows for other options that will not be used here). The number of spins in the extra- and intracellular domains was counted and the fractional populations calculated.

After distributing the spheres and spins in the domain, iterations begin – each representing a step of infinitesimal duration d*t*. We are iterating a single spin at a time, to allow for the use of the parfor function (parallel simulations of multiple spins on multiple processing units). In each iteration step, two spherical angles are drawn from a uniform distribution. The spin is then moved in that direction in 3D a distance 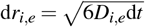, where the choice of *D*_*i*_ or *D*_*e*_ depends on the position of the spin (intra- or extracellular space). We define an indicator variable which holds the values 1 or 2 when the spin is in the intra- and extracellular domains, respectively. If the spin is in a certain sphere, we have a sphere tracker variable that indicates the number of the sphere.

At each step, the code checks whether the spin attempts to cross a membrane. If so, we calculate the crossing probability [89]

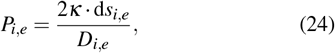

where d*s* is the distance of the spins from the closest point on the sphere from which it exists/enters (here we are actually approximating the curvature of the membrane as flat, assuming d*t* is small enough). A randomly generated number between 0 and 1 then determines whether the particle crosses or not. Note that we use the low-permeability limit relation [44, 90, 91]

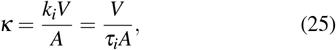

where *V* and *A* are the compartment volume and surface area, respectively. This means that for a spherical compartment of diameter *a*

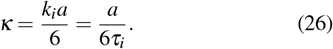

The code differs whether we simulate a CG or a FEXSY experiment. We follow the position of all spins and tag it whenever a gradient is applied, assuming the SGP approximation and using Eqs. (1) and (2). At the measurement time we use a discretized version of Eq. (3)

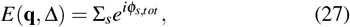

where the summation is over all spins *s*. We then analyze the attenuation curves as if they were attained in an experiment and compare the *τ*_*i*_ that we set as a simulation parameter, and the one inferred using in the analysis of the attenuation curves.

## 3. RESULTS

Here we decided to study exchange in samples of increasing complexity with two different diffusion NMR-based methods. First, yeast cells were studied both before and after a short fixation of 15 minutes (*n* = 3). The more complex optic nerves were then measured before and after 24 hours of fixation (*n* = 3). In each sample, CG-PGSTE and FEXSY experiments were conducted back-to-back to allow for direct comparison.

### A. Exchange in yeast cells

#### i. CG-PGSTE NMR experiments

In Figure 2(a), we plot the decay of the normalized signal *E* vs. the diffusion time Δ in CG-PGSTE NMR experiments, performed on an exemplary yeast sample. In Figure 2(b), we plot instead the natural logarithm ln(*E*). At low q-values, the attenuation is clearly not mono-exponential. However, it is immediately apparent from Figure 2(b) that at high q-values, the faster diffusing population is largely attenuated, even at relatively short diffusion times. This results in attenuation that is, to a good approximation, a mono-exponential attenuation for the slow diffusing population. In fact, a gradient strength of 30 G/cm, with *δ*_*e f f*_ of only 2 ms, which is now available in some clinical MRI scanners [92], is enough to effectively attenuate the signal from the fast population in this sample. Note that at very short diffusion times and high *q* values, we observe a non-monotonic behavior. This does not affect the long time asymptotics and our analysis. Such behavior was not observed in any other sample, and is likely to be related to some fermentation featured by the non-fixed yeast cells.

**FIG. 2.**
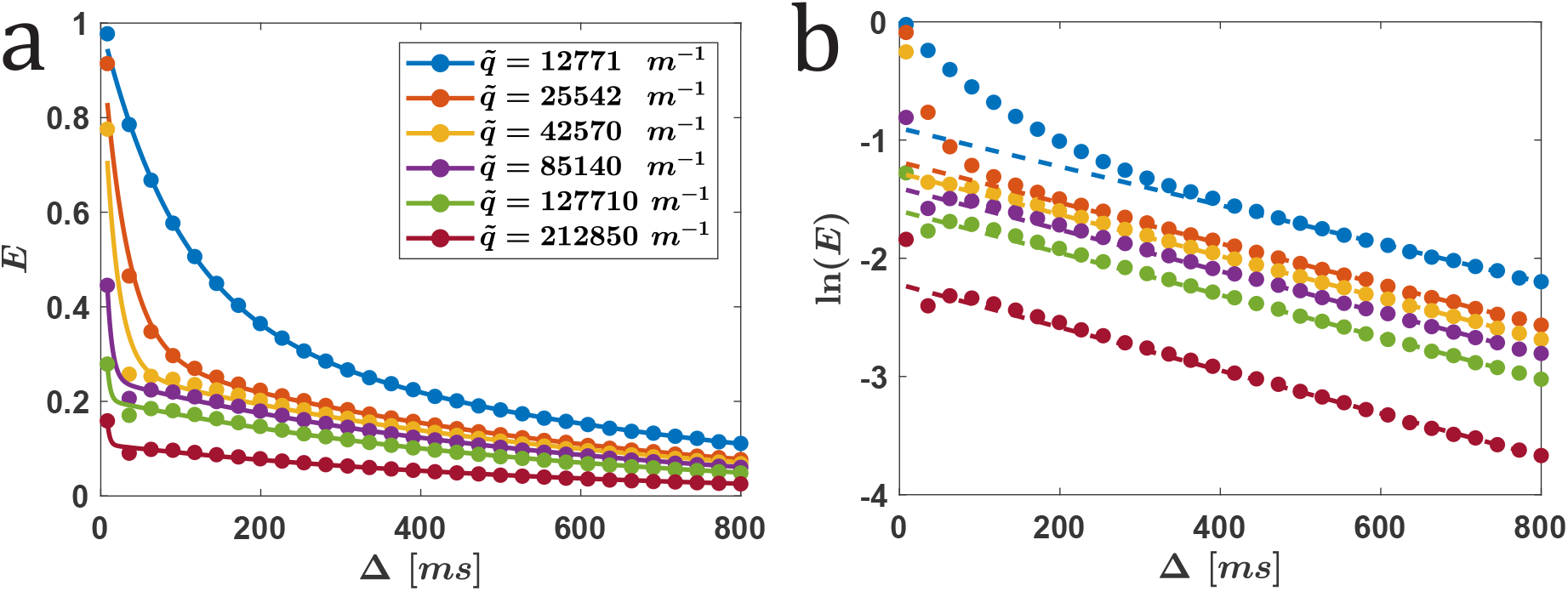
CG-PGSTE NMR experiments on yeast cells: (a) Normalized signal *E* vs. the diffusion time (full circles). Data was fitted with Eq. (6) (lines), and (b) ln(*E*) vs. the diffusion time (full circles). Data was fitted with Eq. (16) using the 10 data points with the highest diffusion times (dashed lines). Data is shown for different q-values obtained with *δ*_*eff*_ = 2 ms and g-values of 15 (blue), 30, 50, 100, 150 and 250 G/cm.

The fit in Figure 2(a) is according to Eqs. (6) and (15) originating from the Kärger bi-compartmental model [26–29]. The fit in Figure 2(b) is the log-linear fit presented in Eq. (16), as suggested by Meir et al. [37, 38]. The intracellular mean residence time (*τ*_*i*_, i.e., 1*/k*_*i*_), was calculated from the fits assuming the slow exchange time limit. The results for an exemplary sample are summarized in Table I and show the dependence of the calculated values on the gradient strength for both analysis methods.

**TABLE I.** The extracted *τ*_*i*_ and *C*_*A*_ (Eq. (6)) values along with *R*^2^ of the fits for an exemplary yeast cells sample. Values were obtained from two different fitting procedures (Eqs. (6) and (15)) of the CG-PGSTE NMR data and are presented as function of the gradient strength (*g*) used.

| Gradient Strength<br>(G/cm) | Bi-exponential fit |  |  | Log-linear fit |  |
| --- | --- | --- | --- | --- | --- |
| | $\tau_i$ (ms) | $C_A$ | $R^2$ | $\tau_i$ (ms) | $R^2$ |
| 15 | 621 | 0.40 | 0.9992 | 613 | 0.9981 |
| 30 | 583 | 0.30 | 0.9883 | 580 | 0.9985 |
| 50 | 602 | 0.27 | 0.9755 | 569 | 0.9995 |
| 100 | 572 | 0.25 | 0.9934 | 569 | 0.9991 |
| 150 | 564 | 0.20 | 0.9924 | 561 | 0.9992 |
| 250 | 555 | 0.11 | 0.9921 | 550 | 0.9976 |

The data presented in Table I clearly show that even for the lower q-value, where the linear regime appears only at relatively long diffusion times, the extracted value for *τ*_*i*_ is only 10-15% off from the one extracted from the highest q-value. When g-values in the range of 30 − 250 G/cm were used (a very short *δ*_*e f f*_ was kept), the extracted *τ*_*i*_ values decreased, in the two types of analysis methods, by only ∼5% and good agreement was observed between the absolute values extracted from both analysis methods. For this range of g-values the average *τ*_*i*_ values extracted from the bicompartmental model and the log-linear fit were found to be 550 ± 36 ms and 574 ± 6 ms, respectively. These results show that, as long as the diffusion weighting is high enough to suppress the fast-diffusing population, the two simplified analysis methods provide very similar and robust results.

The next step was to check for the reproducibility of the CG-PGSTE NMR results. We examined the reproducibility between different yeast cell samples. Three yeast cell samples were prepared and measured using a constant gradient strength of 100 G/cm. Using a log-linear fit, the average *τ*_*i*_ value was found to be 574 ± 6 ms. In addition, we checked the reproducibility within a sample for a period of about 16 hours: One of the samples was measured four times in 4 hour intervals and the average *τ*_*i*_ values extracted from the bi-compartmental model and the log-linear fit were found to be 557 ± 23 ms and 569 ± 11 ms, respectively, showing the repeatability of the extracted *τ*_*i*_ values with time.

After evaluating the reproducibility and repeatability of the obtained results, we used the same methodology to study the effect of formalin fixation on transmembrane exchange in yeast cell samples. Figure S1 shows the same type of data as in Figure 2 for yeast cells after 15 minutes of fixation, and the results of the analysis are summarized in Table II.

**TABLE II.** Same as in Table I but for an exemplary *fixed* yeast cells sample. Note that very high gradients produce poor bi-exponential fits, as the extracellular domain is attenuated even at short diffusion times (see Figure S1).

| Gradient Strength<br>(G/cm) | Bi-exponential fit |  |  | Log-linear fit |  |
| --- | --- | --- | --- | --- | --- |
| | $\tau_i$ (ms) | $C_A$ | $R^2$ | $\tau_i$ (ms) | $R^2$ |
| 15 | 278 | 0.3154 | 0.9998 | 264 | 0.9987 |
| 30 | 286 | 0.0950 | 0.9940 | 311 | 0.9951 |
| 50 | 310 | 0.0691 | 0.9887 | 332 | 0.9940 |
| 100 | 220 | 0.0861 | 0.9988 | 336 | 0.9809 |
| 150 | 110 | 0.1409 | 0.8626 | 321 | 0.9813 |
| 250 | 133 | 0.0762 | 0.8893 | 379 | 0.9841 |

Figure S1 shows that, unlike the non-fixed yeast cell samples, the attenuation of the signal as a function of the diffusion time Δ for yeast cells after fixation does not appear monoexponential even at the highest q-values used in this study. However, on a log-scale, it appears nearly linear for high qvalues and long Δs (Figure S1). All these observations suggest that exchange rates in the fixed yeast cell samples are significantly higher than those in the non-fixed yeast cell samples. Therefore, the validity of Eqs. (15) and (16) may, justifiably, be questioned in this case. However, similar to living cells, the dependence of *D*_*A*_ on 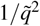 for this sample was linear, albeit with a slightly lower *R*^2^ value, as depicted in Figure 3.

**FIG. 3.**
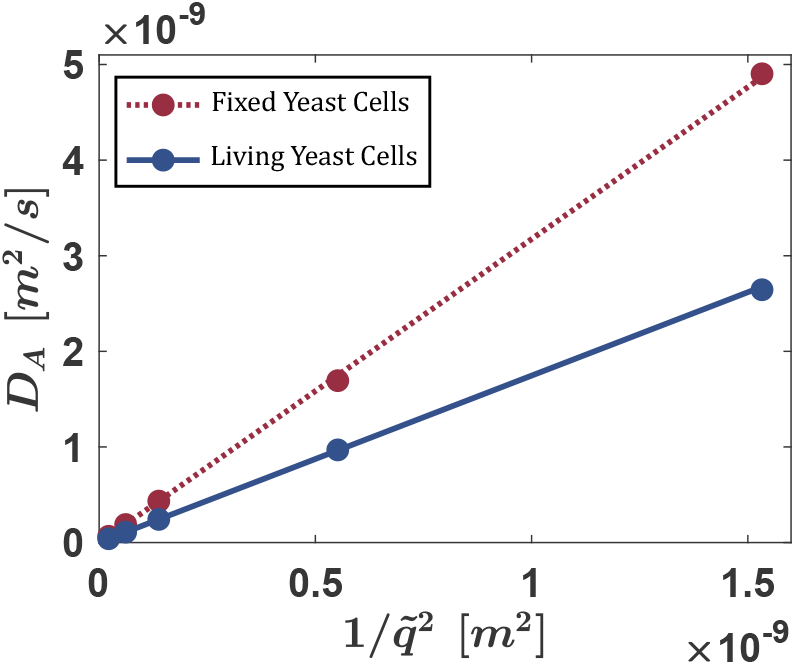
*D*_*A*_ vs. 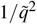 (full circles) along with a linear fit for living yeast cells (solid line with slope corresponding to *τ*_*i*_ of 580 ms, *R*^2^ = 1), and for yeast cells after fixation (dashed line with slope corresponding to *τ*_*i*_ of 316 ms, *R*^2^ = 0.9997).

Table II shows that the log-linear analysis is again reproducible above 30 G/cm, providing an average mean residence time (*τ*_*i*_) of 334 ± 27 ms. For gradient strength in the range of 30 − 250 G/cm the values appear nearly constant; however, it should be noted that the quality of the fits is lower as compared to the living yeast cell sample. The bi-exponential fit gave similar results when gradient strengths of 30 or 50 G/cm were used; however, at higher g-values the extracellular domain is attenuated even at short diffusion times and smaller *τ*_*i*_ values were extracted as a result of a poorer fit (essentially we are left with a single population and the bi-exponential fit becomes inadequate). This can be visually seen in Figure S1.

To verify the reproducibility of the results obtained also for fixed yeast cells, three different samples were prepared and measured using a CG-PGSTE experiment with a constant gradient strength of 50 G/cm. The log-linear fit analysis was used and the average *τ*_*i*_ was found to be 337 ± 10 ms. One of the samples was measured seven times in 2 hour intervals and an average *τ*_*i*_ value of 351 ± 5 ms was obtained. The short fixation process (15 min) of yeast cells almost doubled the exchange rate – a change that is easily detected by CG-PGSTE.

#### ii. FEXSY experiments

FEXSY results on yeast cells, before and after fixation, are presented in Figures 4 and S2, respectively. Figures 4(a) and S2(a) show the natural logarithm of the normalized signal ln(*E*) as a function of diffusion weighting 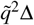 for different mixing times *t*_*m*_. Figures 4(b) and S2(b) present the normalized ADC vs. *t*_*m*_, with fits performed according to the phenomenological Eq. (23). Note that we have used short *δ*, to satisfy the SGP approximation and short TE and diffusion times (8 ms and 13 ms, respectively) to reduce the effect of *T*_2_ relaxation and ensure that exchange occurs mainly during the mixing time. Recall that the FEXSY and CG-PGSTE experiments were conducted consecutively on the same samples, to allow for direct comparison.

**FIG. 4.**
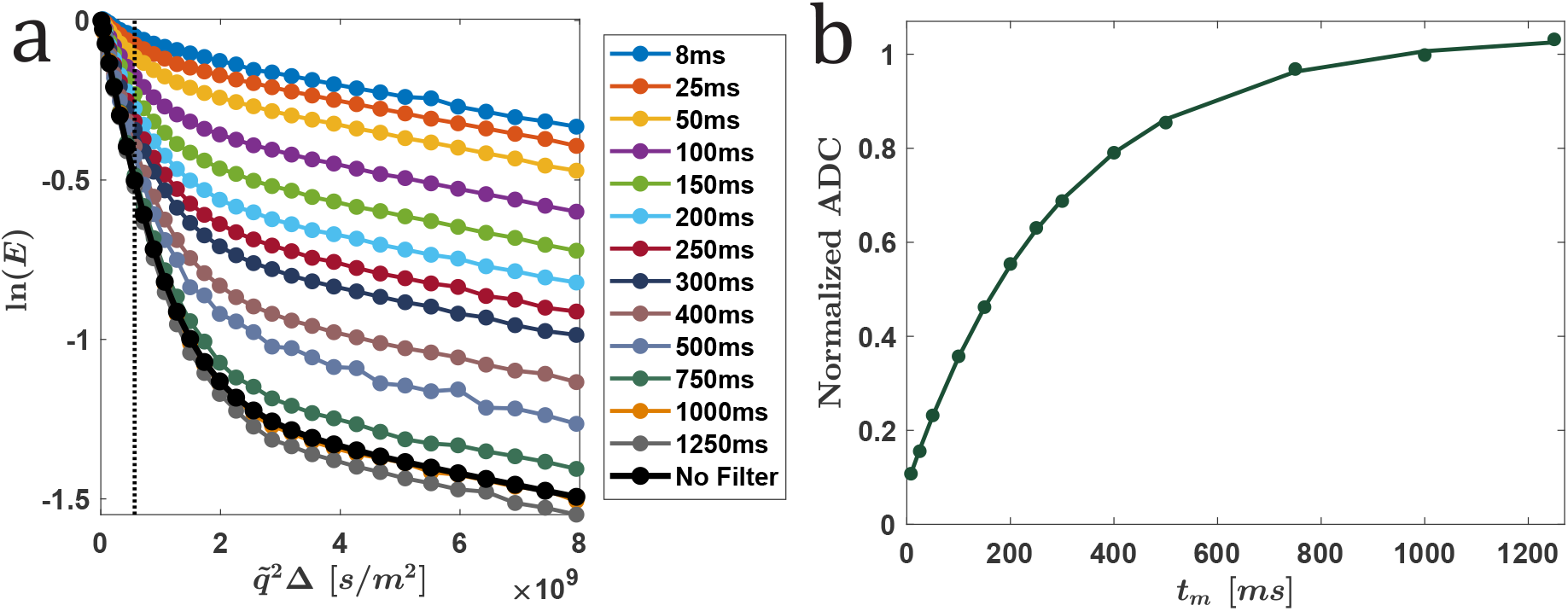
Results of a FEXSY experiment performed on a yeast cell sample: (a) The attenuation of the natural logarithm of the normalized echo ln(*E*) vs. 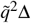 for different *t*_*m*_ values of the FEXSY and the no-filter experiment (full circles), and (b) normalized ADCs extracted (full circles) from the data presented in (a). Note that the gradient strength of the filter was 150 G/cm and that the gradients in the measurement block were incremented from 2.5 to 150 G/cm in 30 steps. The effective *δ* was 2 ms. Normalized ADCs were computed from ADC extracted from the first 8 points of each graph in (a) divided by the ADC extracted from the no filter experiment (ADC_*eq*_). The lines in (a) are just to guide the eyes while the line in (b) is a fit to Eq. (23).

From the data shown in Figures 4(b) and S2(b), using the phenomenological fit (Eq. (18)), averaged *τ* values of 289 ± 11 ms and 117 ± 19 ms (*n* = 3) were extracted for fresh and fixed yeast cell samples, respectively. The ADCs were computed from the first 8 points (Figure S2(b)). Note that after normalizing the data with respect to ADC_*eq*_, the exponential growth of the normalized ADC is predicted to plateau at 1 for long *t*_*m*_s. When this value was a free variable of the fit, its mean value (*n* = 3) was indeed 1.03 ± 0.03 and 0.93 ± 0.05 for the fresh and fixed yeast cell samples, respectively.

When the more demanding and instructive bi-exponential exchange model was used, the following parameters were extracted for the three yeast cell samples: *τ* = 251 ± 2 ms, 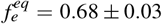, *f*_*e*_ (0) = 0.03 ± 0.01, *D*_*i*_ = 3.65 ± 0.01 · 10^−11^ m^2^*/*s, *D*_*e*_ = 1.37 ± 0.04 · 10^−9^m^2^*/s*. Assuming equilibrium and using Eq. (18), an average *τ*_*i*_ value of 368 14 ms was extracted for these three yeast cell samples. When a sample of yeast cells was measured four times in 4 hours intervals, an average *τ* value of 279 ± 4 ms was extracted using the phenomenological fit and, and of 247 ± 2 ms using the biexponential fit. A similar approach was used to analyze the data from fixed yeast cell samples (*n* = 3), where we used a maximal *t*_*m*_ of 750 ms. There, we obtained the following values: *τ* = 116 ± 15 ms, 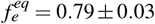, *f*_*e*_ (0) = 0.16 ± 0.05, *D*_*i*_ = 2.55 ± 0.08 · 10^−11^m^2^/s, *D*_*e*_ = 1.34 ± 0.15 · 10^−9^ m^2^*/*s from which an average *τ*_*i*_ value of 146 24 ms was extracted. One fixed yeast cell sample was also measured seven times in 2 hour intervals and there an average *τ*_*i*_-value of 180 ± 23 ms was extracted using the bi-exponential analysis method. Thus, FEXSY detects a significant increase in the exchange rate due to fixation. Note that despite the qualitative agreement, the relative change in the exchange value due to fixation is larger in FEXSY than in CG-PGSTE, suggesting greater sensitivity.

### B. Exchange in optic nerves

After studying exchange in yeast cells both before and after fixation and assessing the reproducibility of the applied methods, we decided to use the same methodologies and analysis to study exchange in optic nerves. Optic nerves, which are more complex systems than yeast cells, were studied before and after 24 hour fixation (*n* = 3).

#### i. CG-PGSTE NMR experiments

Figures 5 and S3 show the results obtained from CG-PGSTE NMR experiments performed on exemplary optic nerves before and after fixation, respectively. These figures show the same information as in Figures 2 and S1, however, since the exchange in the optic nerve is expected to be slower than in yeast cells, here CG-PGSTE NMR experiments were performed with diffusion times (Δs) of up to 1500 ms. In fact, experiments were initially performed with Δs of up to 2000 ms, however, in this range the data quality was low, so we decided not to use it in our quantitative analysis.

**FIG. 5.**
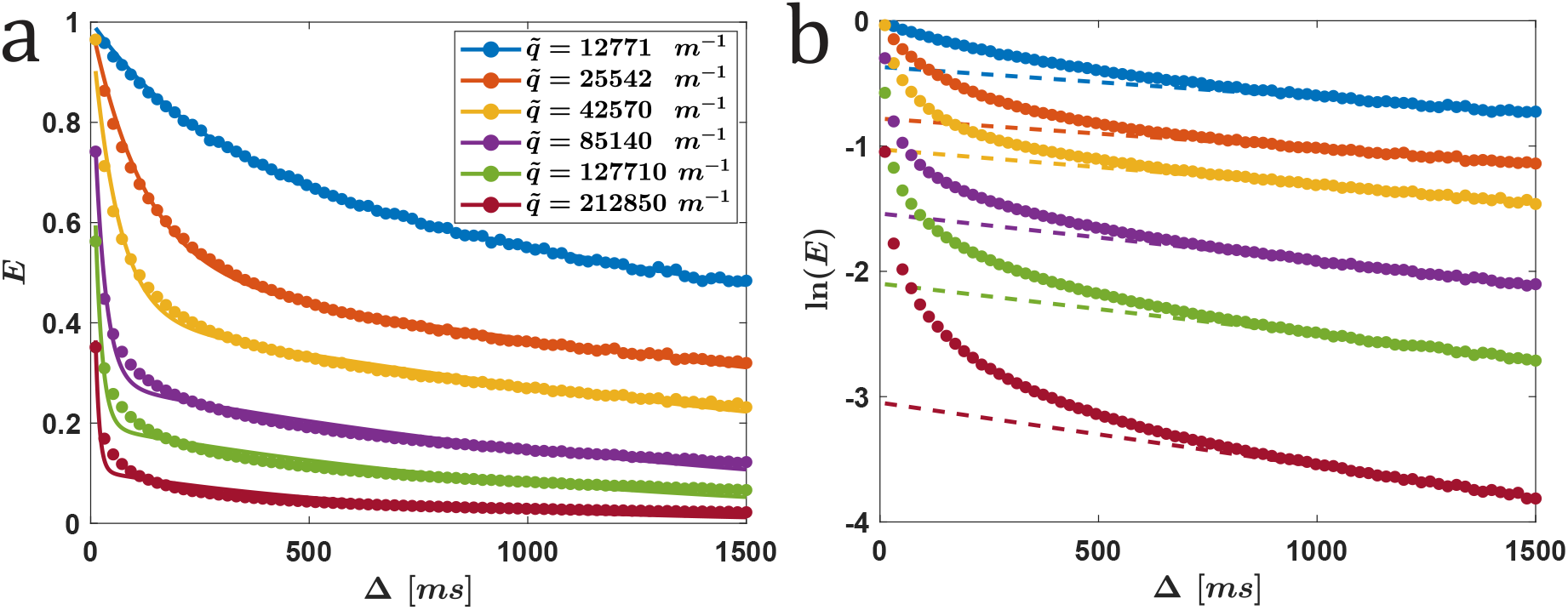
CG-PGSTE NMR experiments on an optic nerve: (a) Normalized signal (*E*) vs. the diffusion time (full circles). Data was fitted with Eq. (6) (lines), and (b) ln(*E*) vs. the diffusion time (full circles). Data was fitted with Eq. (16) using the 20 data points (out of 75 collected) with the highest diffusion times (dashed lines). Data is shown for different q-values obtained with *δ*_*eff*_ = 2 ms and g-values of 15 (blue), 30, 50, 100, 150 and 250 G/cm.

From inspection of the data presented in Figures 5 and S3 it appears that the signal does not attenuate to zero after a long diffusion time even when the diffusion weighting is very high. This may originate from extremely slow exchange or the existence of an additional non-exchanging component that exhibits restricted diffusion. If this is the case, then one should expect the bi-compartmental model to be much less suitable for a quantitative analysis of the data.

For an exemplary sample, the *τ*_*i*_ values before and after formalin fixation, as well as the quality of the fits (*R*^2^), are given in Tables III and IV, respectively. Note that in the case of the bi-exponential fit, the measured values for *τ*_*i*_ do not seem to converge to a constant as the diffusion weighting increases. Clearly, the agreement between the fits and the experimental data for the optic nerve is significantly lower than in the yeast cells, as manifested by the significantly lower *R*^2^ values. In fact, as the gradient strength varies, the extracted *τ*_*i*_ values decrease substantially. The significant dependence on the experimental parameters implies that comparing two reported values is problematic, unless the experiments were performed with the exact same parameters.

**TABLE III.** The extracted *τ*_*i*_ and *C*_*A*_ values along with the *R*^2^ of the fits for the two analysis methods (Eqs. (6) and (15)) of the CG-PGSTE NMR experiments done on an exemplary optic nerve sample. Results are given as function of the gradient strength (*g*) used.

| Gradient Strength<br>(G/cm) | Bi-exponential fit |  |  | Log-linear fit |  |
| --- | --- | --- | --- | --- | --- |
| | $\tau_i$ (ms) | $C_A$ | $R^2$ | $\tau_i$ (ms) | $R^2$ |
| 15 | 3618 | 0.72 | 0.9969 | 4128 | 0.9315 |
| 30 | 3083 | 0.51 | 0.9925 | 4166 | 0.9376 |
| 50 | 2429 | 0.41 | 0.9890 | 3511 | 0.9352 |
| 100 | 1567 | 0.28 | 0.9827 | 2617 | 0.9776 |
| 150 | 1148 | 0.19 | 0.9673 | 2459 | 0.9763 |
| 250 | 692 | 0.11 | 0.9470 | 1977 | 0.9791 |

In the optic nerve case, it can be seen that the dependence of *D*_*A*_ on 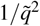 approaches linearity only at very high diffusion weighting, as shown in Figure 6. When the diffusing weighting is increased, by doubling *δ* for example, linearity is observed for higher 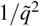 values. This observation is even more apparent in Table S1 which presents the extracted *τ*_*i*_ values for CG-PGSTE experiments conducted with even larger *δ*.

**FIG. 6.**
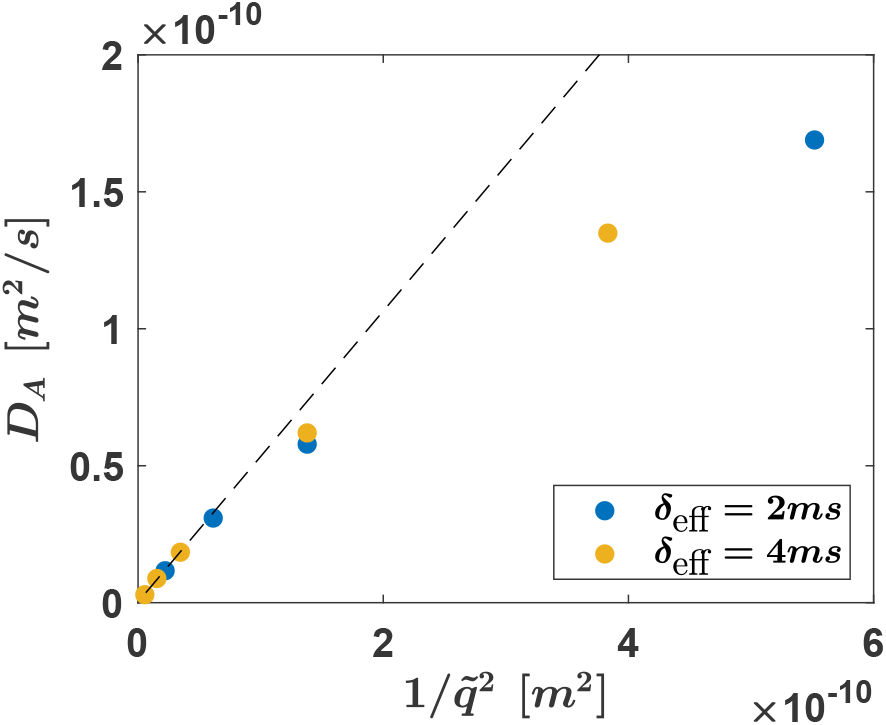
*D*_*A*_ vs. 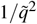 (full circles) for the CG-PGSTE experiments conducted on the optic nerve. Data was collected with *δ*_*eff*_ = 2 and 4 ms. The dashed lines are linear fits to the first two or three data points for *δ*_*eff*_ = 2 and 4 ms, respectively. Their slopes correspond to *τ*_*i*_ values of 1857 and 1759 ms, respectively.

From the analysis of the CG-PGSTE data (*g* = 150 G/cm) of three different optic nerves before and after formalin fixation, averaged *τ*_*i*_ values of 2260 ± 69 ms and 2490 ± 223 ms were extracted, respectively. These results apparently show, as one could have expected, that exchange is significantly slower in myelinated nerves compared to yeast cells. However, we have seen that the bi-compartmental model breaks down in the analysis of the optic nerve, and we can obtain almost any value by changing the experimental parameters, and so one should be cautious before making such a claim. In Sec. 4 we address this issue in detail and suggest a tri-compartmental model that fits the data significantly better.

Although fixation has a clear effect on the permeability of yeast cells, it does not appear to significantly affect the optic nerve. In another set of optic nerves (data not shown), we have seen an increase in exchange of about ∼ 15%, still much lower than the increase of ∼ 66% that was observed in yeast.

#### ii. FEXSY experiments

Figures 7 and S4 show the results of FEXSY experiments carried out on exemplary fresh and formalin-fixed optic nerves, respectively.

**FIG. 7.**
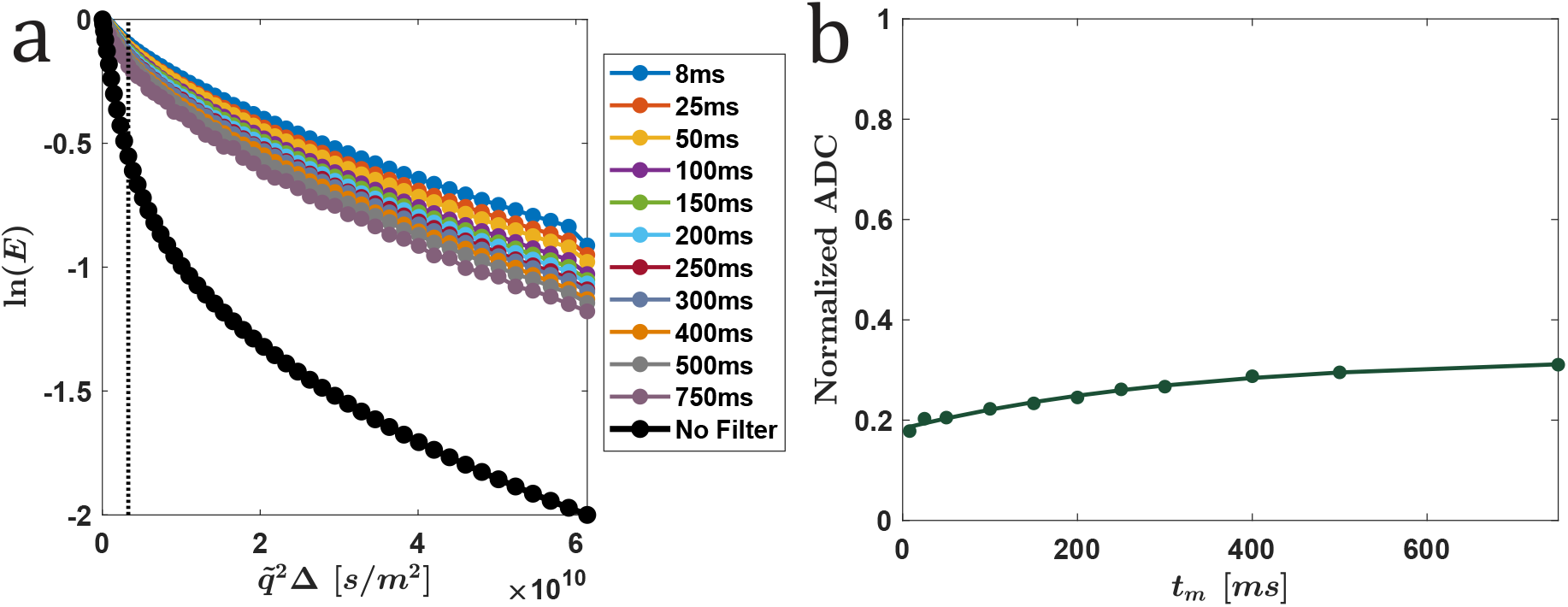
Results of a FEXSY experiment performed on an optic nerve sample: (a) The natural logarithm of the normalized echo ln(*E*) vs. 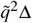 for different *t*_*m*_ values of the FEXSY and the non-filter experiments (full circles), and (b) normalized ADCs extracted (filled circle) from the data presented in (a). Note that the gradient strength of the filter was 250 G/cm and that the gradients in the measurement block were incremented from 2.5 to 250 G/cm in 52 steps. The length of the sine-shaped gradient pulses was 3.14 ms, i.e., *δ*_*eff*_ = 2 ms. For both filter and measurement PGSTE blocks Δ/*δ*_*eff*_ of 35*/*2 ms were used and *t*_*m*_ was varied between 8 and 1000 ms. Normalized ADCs were computed from ADCs extracted from the first 12 points of each graph in (a) divided by the ADC extracted from the no filter experiment (ADC_*eq*_). The lines in (a) are just to guide the eyes while the line in (b), is a fit to the phenomenological model (Eq. (23)).

The data presented in Figure 7(a) shows that indeed even at 750 ms the signal decay does not reach that of the no-filter experiment. Unlike the results observed for yeast cells, the optic nerve data presented in Figure 7(b) shows that the normalized ADC does not approach the asymptotic value of unity. These observations seem to suggest that some of the basic assumptions on which the FEXSY analysis is based do not hold when dealing with optic nerves. Nevertheless, for completeness, we performed the analysis (the same as in Sec. 3 A ii).

Limiting our analysis to a maximal *t*_*m*_ of 750 ms, the phenomenological fit in Figure 7(b) gives an average *τ* value of 348 ± 108 ms (*n* = 3). Importantly, the plateau value was a free variable of the fit, and its value 0.37 ± 0.07, very far from unity. When constraining the fit to plateau at 1.00, as the model suggests, the resulting *τ* value is 3187 ± 1029 ms – an order of magnitude larger! Under this constraint, the fit is visibly poor (and *R*^2^ = 0.92 ± 0.03). When surveying the literature (for example, see the plots in Ref. [50]) it seems that such a constraint might have been employed in the analysis, leading to a significantly smaller and inaccurate AXR.

The bi-exponential global fit provided the following values (*n* = 3): *τ* = 254 ± 34 ms, 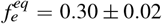, *f*_*e*_ (0) = 0.12 ± 0.01, *D*_*i*_ = 1.22 ± 0.12 · 10^−11^ m^2^*/*s, *D*_*e*_ = 1.78 ± 0.21 · 10^−10^ m^2^*/*s. We thus have 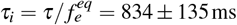. Note that the *τ* values extracted for the optic nerves are very similar to those measured in yeast. After dividing by the extracellular fraction, we get a significantly larger *τ*_*i*_ in the optic nerves, since 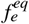 is much smaller. However, we have just observed that the signal does not relax to unity and the picture of two exchanging fractions is inadequate. Therefore, in optic nerves, reporting *τ*_*i*_ obtained in this way appears to be incorrect. In Sec. 4 we address this issue in detail and suggest a tri-compartmental model that fits the data significantly better.

When the same analysis was performed on the optic nerves after 24 hours of fixation, we obtained *τ* = 220 ± 64 ms,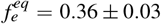, *f* _*e*_ (0) = 0.15 ± 0.01, *D*_i_ = 1.14 ± 0.13 · 10^−11^ m^2^*/*s, *D*_*e*_ = 1.91 ± 0.09 · 10^−10^ m^2^*/*s. We thus have 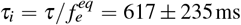. Comparing the FEXSY results before and after fixation, we see an increase in the exchange rate of about ∼ 33%, which is much lower than the ∼ 150% increase observed in fixed yeast cells, albeit still detectable.

### C. Simulations

We first challenged our algorithm with the two well-known limiting cases: (i) Free diffusion, where the attenuation curve is given by Eq. (5); and (ii) Diffusion restricted to a spherical compartment, where the attenuation curve is given by Eq. (9). These two cases correspond to *κ* → ∞ and *κ* → 0, respectively. In Figure S5 we show that the simulation results for these two cases match the analytical formulas.

We then proceeded to simulate a domain with homogeneous permeable spheres, namely with a finite *κ*, that corresponds to a known *τ*_*i*_ = 550 ms according to Eq. (26). This system is an idealization of the yeast cell sample. We tested the system with random packing of the spheres of moderate density, but also tested two limiting cases, that of an HCP lattice and that of a lower sphere density (see Figure S6). In both limiting cases, one might expect the model to break down. When the sphere density is very low, the validity of the well-mixing assumption might be questioned (as the spins can wonder off to infinity, being completely free and unrestricted in the extracellular space). In the opposite scenario, the HCP lattice is a close packing arrangement, where the modified Kärger model is expected to break down due to the high tortuosity of the extracellular space.

For simplicity from here onward we will denote the HCP lattice as set-up (i), the 43 randomly packed spheres as set-up (ii), and the 23 randomly packed spheres as set-up (iii).

We started by simulating a CG-PFG experiment with *g* = 100 G/cm and *δ*_*e f f*_ = 2 ms (hence 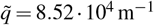), and employing the SGP approximation. The results were fitted ac-cording to Eq. (16), and are presented in Figure 8. Using the log-linear fit we obtain *τ*_*i*_ of (i) 611 ms, (ii) 565 ms and (iii) 623 ms. The errors are less than 15% even for the most extreme case of an HCP lattice, and for a moderate density we attain an error of only 3%. In the same order, we obtain 589 ms, 532 ms and 515 ms using the bi-exponential fit.

**FIG. 8.**
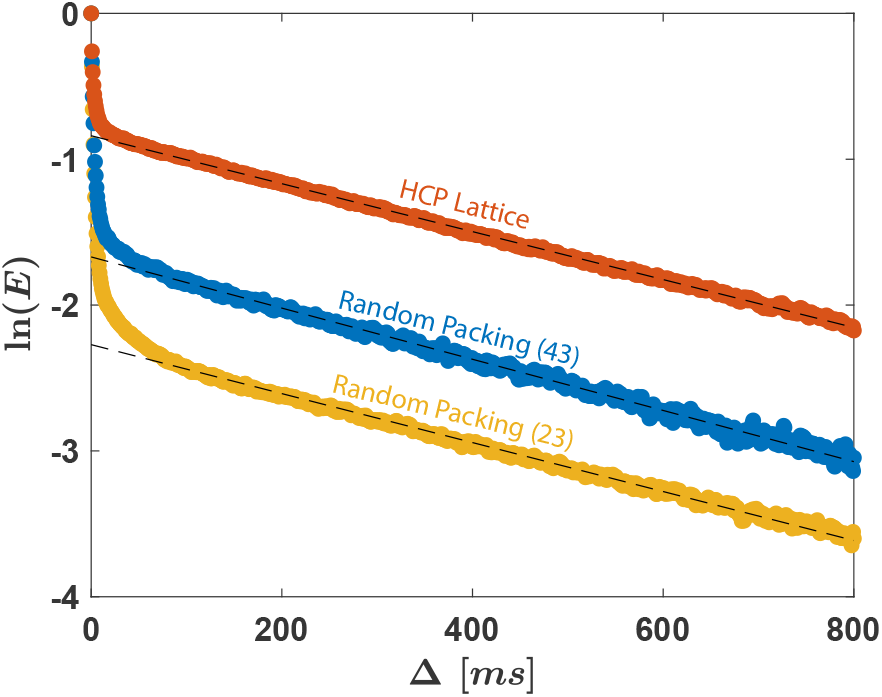
Monte-Carlo simulations of a CG-PFG experiment with 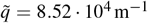. Water molecules (spins) propagating in a domain containing homogeneous permeable spherical cells with radius of 5*µ*m and permeability *κ* = 1.515 · 10^−3^ *µ*m/ms. According to Eq. (26), this value of permeability corresponds to a mean residence time of *τ*_*i*_ = 550 ms. The upper dots (red) correspond to an HCP lattice and the middle (blue) and bottom (yellow) dots correspond to a random packing of 43 and 23 spheres, respectively. The spins were initially uniformly distributed inside a 22x22x22 *µm*^3^ box (see Figure S6). We simulated 10^5^ spins with d*t* = 1 *µ*s, where we set *D*_*e*_ = *D*_*i*_ = 2 *µ*m^2^*/*ms. In the cases of random packing, the attenuation of the signal is greater, due the free diffusion in the extracellular space, and so 10^6^ were simulated. The dashed lines are fits according to Eq. (16). The *τ*_*i*_ extracted using the log-linear fit (using the known 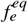), from top to bottom, are 611 ms, 565 ms, 623 ms. It can be appreciated that these values are close to the in-silico value of 550 ms, with an error of less than 15% even for the limiting cases. For the moderate density where the bi-compartmental model is expected to work the best, 43 spheres with 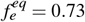, the error is only ∼ 3%.

On the same three packing arrangements we have simulated FEXSY experiments, with the same parameters as used in the yeast cells experiments. Using the phenomenological fit we obtained *τ* values of (i) 231 ms, (ii) 442 ms and (iii) 535 ms. Unlike the situation in real experiments, in silico we know the equilibrium extracellular fractions, so we can esti-mate how accurate these *τ*s are. Here, they are (i)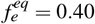, (ii) 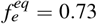 and (iii) 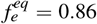, respectively (where for the HCP, the overall density is higher than the local density of 0.26 due to the finiteness of the domain and contributions from the boundaries). Using these, we obtain *τ*_*i*_ values of (i) 578 ms, (ii) 604 ms and (iii) 625 ms.

Using the bi-exponential fit we obtain: (i)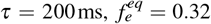, hence *τ* = 623 ms; (ii)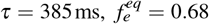, hence *τ*_*i*_ = 566 ms; (iii)τ = 488ms, 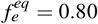, hence *τ*_*i*_ = 606 ms. Note the underestimation of 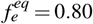 by ∼7 − 20%, since *τ* values are also underestimated, taking the ratio cancels the errors such that the estimation of *τ*_*i*_ can be more accurate. As in the case of CG-PFG, the errors for the limiting case are surprisingly small, and the best inference is for the case of moderate sphere density, with an error of only ∼ 3%.

Note that for the simulated data, just as for the experimen-tal data, *τ* calculated from the phenomenological fit is higher than *τ* calculated from the bi-exponential global fit. However, now we can observe that the latter analysis tends to be more accurate. Thus, if one collects the entire FEXSY data anyway, it is better to use the global fit (which also provides more information). If one wishes to shorten the measurement time, *τ* and 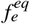 can be attained by using CG-FEXSY [48].

In Figure 9, we show the data for the HCP lattice. We chose to show this data because it behaves qualitatively as the bi-compartmental model predicts (the signal relaxes back to unity). This is despite the high density of cells and the fact that the extracellular space is even more restricted than the intracellular space. This result is important: We can now rule out high density and restricted extracellular domain as reasons for the breakdown of the model when analyzing WM data (recall that in Figure (7) the signal does not relax to unity, leading to erroneous measurement of the exchange rate if one forces this in the fit).

**FIG. 9.**
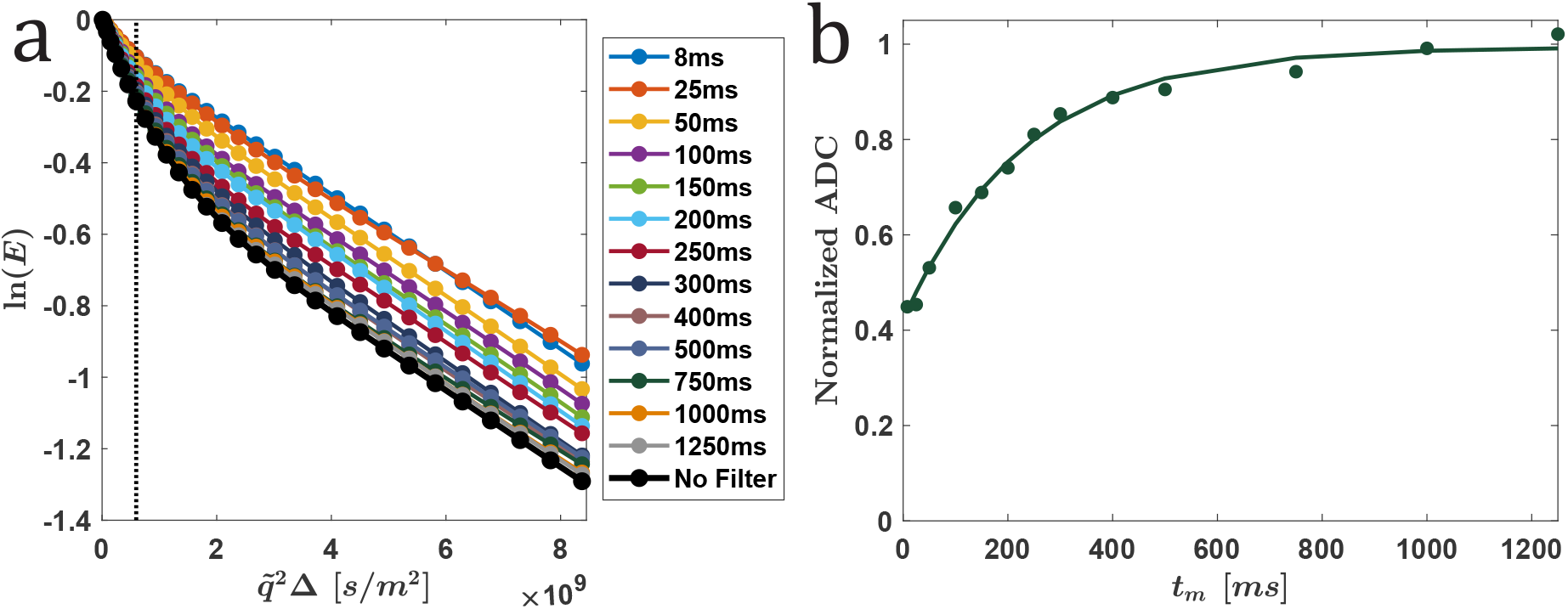
Monte-Carlo simulations of a FEXSY experiment on a HCP lattice. Water molecules (spins) propagating in a domain containing homogeneous permeable spherical cells with radius of 5 *µ*m and permeability *κ* = 1.515 · 10^−3^ *µ*m/ms. According to Eq. (26), this value of permeability corresponds to a mean residence time of *τ*_*i*_ = 550 ms. The NMR parameters are the same as the experimental parameters in Figure 4. We simulated 10^5^ spins with d*t* = 1 *µ*s, where we set *D*_*e*_ = *D*_*i*_ = 2 *µ*m^2^*/*ms. (a) The attenuation of the natural logarithm of the normalized echo ln(*E*) vs. 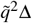 for different *t*_*m*_ values of the FEXSY and the no-filter experiment (full circles), and (b) Normalized ADCs were computed from the first 10 points of each curve in (a), divided by the ADC extracted from the no-filter experiment (ADC_*eq*_). The lines in (a) are just to guide the eyes while the lines in (b) are fits to Eq. (23), and we obtain *τ* = 231 ms. Using a bi-exponential fit of the data (a) we compute 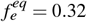, hence *τ* = 623 ms. Since the HCP is a limit case where we would expect the model to break down, an error of 13% is a surprisingly good result. Importantly, note that despite the high density of the cells and the fact that the extracellular space is of high tortuosity, the signal relaxes back to unity. This is in sharp contrast with results from the optic nerve in Figure 7, thus ruling out high cell density as an explanation for why the model breaks down in the case of the optic nerve.

All of the simulations above were performed with *D*_*e*_ = *D*_*i*_ = 2 *µ*m^2^*/*ms. We have kept the ratio of the intrinsic diffusion coefficients and changed the absolute value to *D*_*e*_ = *D*_*i*_ = 1 *µ*m^2^*/*ms. No significant effect. We then changed the ratio. When *D*_*e*_ *> D*_*i*_, we did not observe any significant error. However, when *D*_*i*_ = 2 *µ*m^2^*/*ms and *D*_*e*_ = 1*µ*m^2^/ms we extracted faster apparent exchange rate with an error of ∼ 35% and when *D*_*i*_ = 2 *µ*m^2^*/*ms and *D*_*e*_ = 0.5 *µ*m^2^*/*ms we observed an error of ∼ 70%. Both methods gave very similar errors.

Lastly, in Sec. 1 we mentioned by passing that exchange, if not taken into account in the model, can lead to an error in the determination of, for example, the compartment size or the relative fractions of spins in the two domains. In Appendix B, we demonstrate this effect with a set of simulations of cells with increasing exchange, and the attempt to analyze the qspace data via a simple bi-compartmental model that assumes no exchange. Interestingly, we observe high robustness of the extracted cell radius, and to a lesser extent the fractional populations, with significant error starting to show only at relatively high exchange rates, ∼ 10 − 1000 s^−1^ and higher. Thus, only the highest exchange rates measured here, those of fixed yeast cells (∼ 10 s^−1^) raise a concern for a small error if exchange is neglected. However, exchange values that are larger than 100 s^−1^ were recently reported in the literature, see discussion in Sec. 6 H.

## 4. TOWARDS CORRECT MODELING OF EXCHANGE IN WHITE MATTER

In Sec. 3 B we observed that both methods fail to describe the optic nerve signal due to inadequate modeling. In Sec. 3 C we have used simulations to narrow down the scope of possible explanations. The explanation that prevailed and seems most likely is that in white matter there is a violation of the premise of the Kärger model, which assumes the existence of only two exchanging populations. We may instead be dealing with a tri- or multi-compartmental system, in which at least some compartments do not exchange with one another.

The results suggest that a tri-compartmental model will do, albeit a more elaborate multi-compartmental system cannot be dismissed. Nonetheless, let us begin by deriving a tricompartmental model that will fit the data: Two exchanging compartments and an additional compartment that does not exchange with any other. We will first assume that the spins in the additional compartment are freely diffusing, and then impose a restriction to test if the results change.

### A. Tri-compartmental model for CG-PFG experiments

Adding a third compartment, which is non-exchanging, Eq. (10) becomes

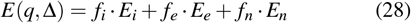

where the subscript *n* denotes the non-exchanging compartment and we have 1 = *f*_*i*_ + *f*_*e*_ + *f*_*n*_.

If one assumes that all compartments are freely diffusing, Eq. (11) is replaced by

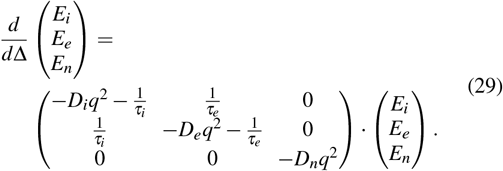

We can transform the matrix to an uncoupled coordinate system and solve to give a tri-exponential expression with the form of Eq. (6), but with effective diffusion coefficients (*D*_1_, *D*_2_ and *D*_3_) and effective fractional populations (*C*_1_, *C*_2_, *C*_3_).

Alternatively, we can follow Price and assume that the intracellular compartment is restricted to a sphere. Here again, since the third compartment is not coupled to the other two, the result is similar to the result of the bi-compartmental model with *D*_1,2_ = *D*_*A,B*_ in Eq. (14), *D*_3_ = *D*_*n*_. Similarly, we have *C*_1,2_ = *C*_*A,B*_ and *C*_*n*_ = *f*_*n*_. Thus, we are again dealing with familiar expressions, and as performed for the bicompartmental model, in the slow exchange limit, *q*^2^*D*_*e*_ ≫ *k*_*i*_, *k*_*e*_, we can assume

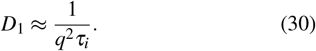

Although it remains generally true that one can attempt to fit the general expression

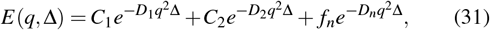

the number of parameters went up from the five of the bicompartmental model to seven. We are thus faced with the question: Is it possible to analyze the long diffusion time asymptotics similarly to the bi-compartmental analysis? Namely, can we still use Eq. (16)?

The answer depends on the value of *D*_*n*_ relative to *D*_1_. In case *D*_*n*_ ≫ *D*_1_, the answer is yes. Given that the difference in the intrinsic diffusion coefficients in vivo is not large, the meaning of this result is that Eq. (16) should work.

However, the above derivation is only correct when the additional non-exchanging population is freely diffusing. If the additional compartment is restricted to, say, a sphere of diameter *a*_*n*_, we obtain

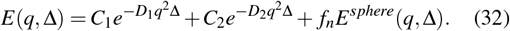

Now, taking the limit *ξ* ≫ 1 we employ Eq. (9) and obtain under the usual assumptions

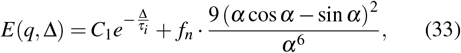

where 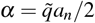.

Importantly, attempting to plug Eq. (33) into the right-hand side of Eq. (16) no more results in the simple left-hand side *τ*_*i*_. Moreover, the result depends on the gradient strength used. In cases where the additional compartment is restricted, Eq. (16) is invalid!

If we are interested solely in measuring the exchange rate, Eq. (33) can be re-written in the simple form

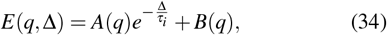

where *A*(*q*) and *B*(*q*) are q-dependent, but are otherwise constant when we perform a CG fit and vary Δ. Note that if Eq. (34) is used analyze a truly bi-compartmental exchanging system, one should get *B*(*q*) = 0 for all *q* values. We verified that this is indeed the case when analyzing the yeast data presented in Sec. 3 A i, in line with our previous conclusion that the signal from yeast samples is well described by the bi-compartmental model.

Re-inspecting the results in 3 B i, we can attribute, at least partially, the dependence of the measured exchange rate on the gradient strength (see tables III and IV) to the existence of a third *restricted* compartment in WM.

**TABLE IV.** Same as in Table III but for an exemplary *fixed* optic nerve sample.

| Gradient Strength<br>(G/cm) | Bi-exponential fit |  |  | Log-linear fit |  |
| --- | --- | --- | --- | --- | --- |
| | $\tau_i$ (ms) | $C_A$ | $R^2$ | $\tau_i$ (ms) | $R^2$ |
| 15 | 3739 | 0.76 | 0.9975 | 3433 | 0.9501 |
| 30 | 2934 | 0.53 | 0.9947 | 3879 | 0.9467 |
| 50 | 2629 | 0.42 | 0.9906 | 3808 | 0.9221 |
| 100 | 1749 | 0.30 | 0.9844 | 2787 | 0.9590 |
| 150 | 1267 | 0.22 | 0.9732 | 2406 | 0.9628 |
| 250 | 753 | 0.12 | 0.9549 | 1905 | 0.9819 |

### B. Tri-compartmental model for FEXSY

Here again we shall first consider that the spins in the third non-exchanging population are freely diffusing. Given that the FEXSY filter module attenuates the signal from this population with a filter efficiency of *σ*_*n*_ = 1, the original global fit analysis should work as is. Put differently, the no-filter experiment is not required for the global fit, where each curve corresponds to a different *t*_*m*_ and normalized according to its first point (*q* = 0 of the measurement block). The additional population is simply filtered out before the mixing time, and because it is not participating in exchange it does not come back and has no effect on the normalized signal.

However, when considering the phenomenological fit of the ADC, caution is needed. There, since the procedure is to normalize according to *ADC*_*no filter*_, the curve will not relax to 1. Practically, the meaning of this result is that instead of forcing the phenomenological fit to relax to the value 1, one should allow a fit with a free parameter *c*:

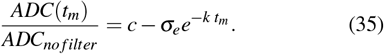

Indeed, in Sec. 3 B ii we have shown that the data does not relax to 1 and that by forcing the fit one introduces an order-of-magnitude error to the measurement of exchange rates. Crucially, as long as *c* is kept as a free parameter of the fit, the extracted *τ* will be accurate. Then, to obtain *τ*_*i*_ one should use the global fit or the CG-FEXSY analysis as before. In fact, *c* can be used to gauge whether the system is bi-compartmental or not.

In many scenarios, the difference in the intrinsic diffusion coefficients of different populations in vivo is small enough such that the above result is expected to be quite accurate, given that the third population is freely diffusing.

However, combining the findings of Secs. (3 B i) and (4 A), we conclude that it is highly likely that the additional compartment in the white matter is restricted. Then, the above model is inadequate, as the third population will not be filtered completely. After applying the filter, we must account for it by writing

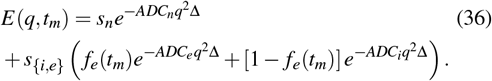

where 0 ≤ *s*_{ *i,e*},*n*_ ≤ 1 are parameters that comprise the filter efficiencies and population size. Since the third compartment is non-exchanging, *s*_*n*_ is determined solely by the filter efficiency and the population size of the third compartment. Similarly, *s* _*i,e*_ groups the signal efficiencies and population size of the two exchanging populations, the intra- and extracellular domains. In this manner, the original definitions of *f*_*e*_(*t*_*m*_) and *f*_*i*_(*t*_*m*_) are retained, and Eq. (18) can be used as before to calculate *k*_*i*_.

### C. Simulations of a Tri-Compartmental System

To corroborate the tri-compartmental model, we simulated set-up (i) from Sec. (3 C), but with an additional nonexchanging compartment in the form of a non-permeable extension of the original domain, see Figure S7. To achieve restriction, the width of the added box was set to 2*µ*m.

Simulating CG-PFG experiments with gradient strengths varying between 15 − 150 G/cm. The results were fitted according to Eq. (34), and are presented in table S2. Averaging over gradient strength we obtain *τ*_*i*_ = 560 ± 44 ms, where the in silico value of 550 ms is in the error range.

For comparison, forcing the bi-exponential fit in Eq. (16) resulted in *τ*_*i*_ = 6855 ± 1008 ms, which is also *q*-dependent (as in Table III). Forcing an incorrect model led to inaccurate and unreliable results.

We then proceeded to simulate a FEXSY experiment, setting *τ*_*i*_ = 550 ms. The results are shown in Figure (10). It can be appreciated that the addition of the third compartment results in a signal attenuation that resembles the data from the optic nerve (Figure 7). Using the tri-compartmental fit in Eq. (36) we compute 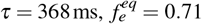, from which we obtain *τ*_*i*_ = 517 ms – very close to the set value. In contrast, forcing the bi-compartmental fit results in the same *τ* but a different 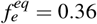, from which we obtain an erroneous value of *τ*_*i*_ = 1018 ms.

We repeated the simulation setting *τ*_*i*_ = 250 ms. Using the tri-compartmental fit in Eq. (36) we compute 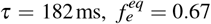, from which we obtain *τ*_i_ = 272 ms. Again, a bicompartmental fit results in an erroneous value of *τ*_*i*_ = 726 ms. Based on these simulations, we conclude that the theoretical model derived herein is indeed adequate for the analysis of tri-compartmental systems. We thus move on to re-analyze the WM data.

**FIG. 10.**
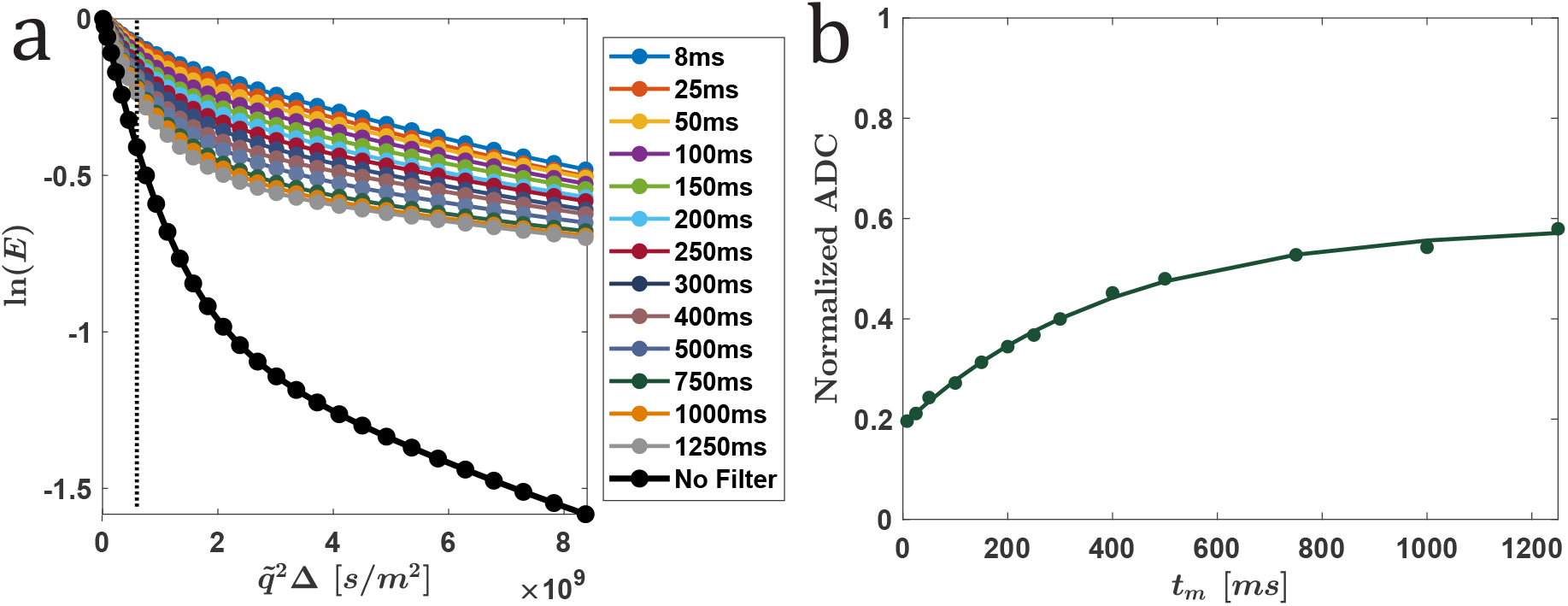
Monte-Carlo simulations of a FEXSY experiment on set-up (i) described in Sec. 3 C, with an additional third restricted and nonexchanging compartment (see Figure (S7)). The additional compartment leads to results which resembles the optic nerve data in Figure (7). In the simulation, we set the mean residence time of *τ*_*i*_ = 550 ms. The NMR parameters are the same as the experimental parameters in Figure 4. We simulated 10^6^ spins with d*t* = 1*µ*s, where we set *D*_*e*_ = *D*_*i*_ = 2 *µ*m^2^*/*ms. (a) The attenuation of the natural logarithm of the normalized echo ln(*E*) vs. 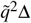 for different *t*_*m*_ values of the FEXSY and the no-filter experiment (full circles), and (b) Normalized ADCs were computed from the first 8 points of each curve in (a), divided by the ADC extracted from the no-filter experiment (ADC_*eq*_). The lines in (a) are just to guide the eyes while the lines in (b) are fits to Eq. (35), and we obtain *τ* = 397 ms. Using a tri-exponential fit of the data (a) according to Eq. (36) we compute 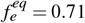, from which we obtain *τ*_*i*_ = 517 ms.

### D. Re-Analyzing WM data using the tri-compartmental model

Recall that in Sec. (3 B) we analyzed optic nerve data while assuming the current bi-compartmental models. Results from using both methods indicate that the bi-compartmental models are inadequate, and that theory and simulations point to the existence of an additional compartment. In this subsection we will put our predictions to the test by re-analyzing the optic nerve data using the tri-compartmental models developed in Secs. 4 A and 4 B.

#### i. CG-PGSTE NMR experiments

In Figure 11 we plot again the same data that was plotted in Figure 5, but now fitted according to Eq. (34) of the tri-compartmental model. This equation was derived for long diffusion times. In practice, we chose to take the 50 highest points, as these produced the highest *R*^2^ values. The results are summarized in Table V.

**FIG. 11.**
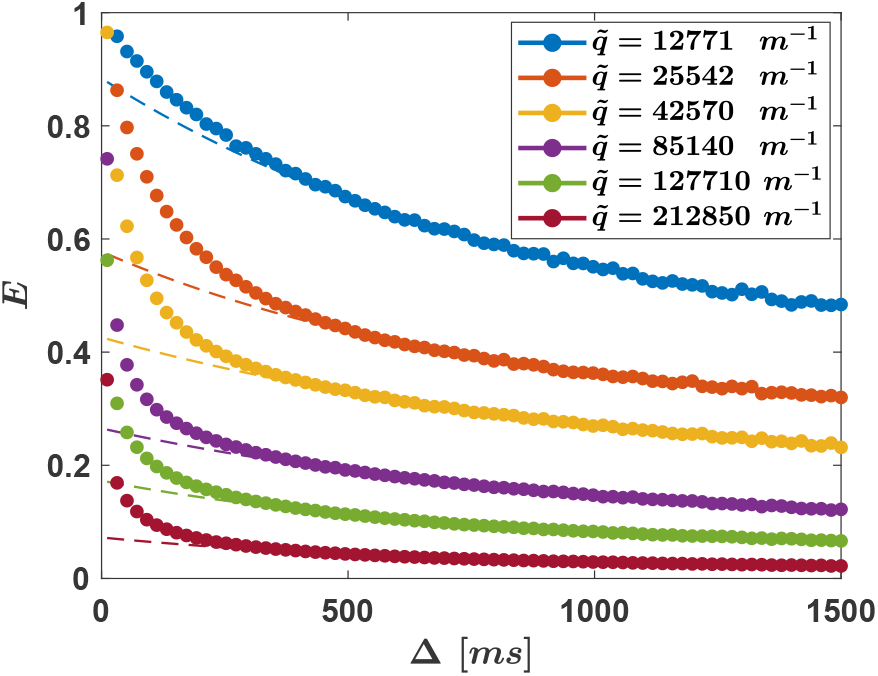
CG-PGSTE NMR experiments on an optic nerve, the same data as in Figure 5(a): Normalized signal (*E*) vs. the diffusion time (full circles). Here, however, data was fitted with Eq. (34) using the 50 data points (out of 75 collected) with the highest diffusion times (dashed lines).

**TABLE V.** Same as in Table III but the fit was performed according to Eq. (34).

| Gradient Strength<br>(G/cm) | Tri-compartmental fit |  |  |  |
| --- | --- | --- | --- | --- |
| | $\tau_i$ (ms) | $A$ | $B$ | $R^2$ |
| 15 | 887 | 0.49 | 0.39 | 0.9964 |
| 30 | 800 | 0.30 | 0.27 | 0.9961 |
| 50 | 1003 | 0.25 | 0.18 | 0.9961 |
| 100 | 922 | 0.18 | 0.09 | 0.9984 |
| 150 | 728 | 0.12 | 0.05 | 0.9986 |
| 250 | 653 | 0.06 | 0.02 | 0.9987 |

First, unlike the case of yeast cells where we get *B*(*q*) ≈ 0 (data not shown), a substantial contribution from *B*(*q*) can be observed. This is yet another indication that the tricompartmental fit is more adequate in describing the WM data.

Importantly, it is readily observed that using the tricompartmental fit the variation of the extracted *τ*_*i*_ as function of the gradient strength is dramatically reduced (recall that the bi-compartmental fit resulted in a range of values that spans an order of magnitude). Looking at an exemplary sample and averaging over all measured gradient strengths, we obtain *τ*_*i*_ = 832 ± 129 ms. We can thus see that with better modeling the results are more robust for change in the experimental parameters.

From the tri-compartmental analysis of the CG-PGSTE data (*g* = 150 G/cm) of three different optic nerves before and after formalin fixation, averaged *τ*_*i*_ values of 730 ± 40 ms and 803 ± 16 ms were extracted, respectively. These values are considerably smaller than the inadequate values calculated in Sec. (3 B i) using the bi-compartmental analysis. The apparent intracellular MRT turns out to be not far from the CGPGSTE measurement of *τ*_*i*_ = 569 ± 11 ms for yeast cells.

#### ii. FEXSY experiments

In Sec. (3 B ii) we already discussed the fact that Figure 7(b) does not plateau at 1. We performed the fit to a function of the same form as in Eq. (35) and obtained a plateau at *c* = 0.37 ± 0.07, and *not* at *c* = 1 as expected for two exchanging compartments. More importantly, when *not* forcing the fit to plateau at 1, we found the average *τ* value of 348 ± 108 ms instead of 3187 ± 1029 ms (*n* = 3).

Performing a global tri-compartmental fit of the data according to Eq. (36) we obtain *τ*_*i*_ = 530 ± 125 ms and *τ*_*i*_ = 387 ± 104 ms (*n* = 3) for the nerves before and after fixation, respectively. Again, we observe that FEXSY is more sensitive to fixation than CG-PFG.

We obtain 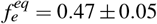 and 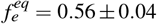 before and after fixation, respectively. Note that while the value of *τ* did not change significantly, the extracted fractional population is dramatically different now that a large portion of the signal is attributed to the third compartment. In turn, the corrected evaluation of *f*_*e*_ leads to a corrected *τ*_*i*_ that is much smaller than the value that we would have reported if forcing a bi-compartmental fit.

## 5. A POTENTIAL EXPLANATION FOR THE QUANTITATIVE DISCREPANCY BETWEEN CG-PFG AND FEXSY

In 2017, Eriksson et al. proposed a model that incorporates spin-spin relaxations into the bi-compartmental model [47]. In order to fit this model to the FEXSY data, an additional complementary data set must be acquired: a series of PGSE based D – *T*_2_ correlation measurements. This experiment is sensitive to the spin-spin relaxation constant *T*_2_, and more specifically, it allows the detection of the difference in intra- and extracellular relaxations, if such exists.

Eriksson et al. have conducted the experiments on a yeast sample. Values of 18 ms and 35 ms were measured for the intra- and extracellular compartments, respectively. They showed that omission of the difference in *T*_2_ values from the model results in a significant error in the measured exchange rate. In their case, the error was ∼ 20%, but it is important to note that the error is highly sample-dependent. First, the intra- and extracellular fractional populations vary between samples. The magnitude of the error is further dependent on the TE used in the FEXSY experiment. Lastly, the *T*_2_ values can vary between different yeast strains and preparation protocols [47].

Here we hypothesize that, compared to CG-PGSTE, FEXSY is much more sensitive to the difference in *T*_2_ values, and is in turn prone to significant errors if the procedure suggested by Eriksson et al. is not performed. To see why, let us revisit the pulse sequences in Figure 1. In the context of our discussion here, the main difference between the two experiments is the way in which the fast population is filtered prior to the measurement. In the FEXSY experiment, the fast extracellular population is filtered out *before* the mixing time and the measurement block. When the measurement block begins, and as we increase *t*_*m*_, the spins are already mixed between the two compartments. Hence, the spins that contribute to the signal are already mixed in environments of different *T*_2_ values. In contrast, the assumption behind the log-linear fit of the CGPGSTE experiment is that the diffusion times are long enough such that the extracellular compartment is consistently attenuated. Hence, the contributions to the signal come mainly from intracellular spins, and spins that succeed in crossing the membrane are rapidly weighted out thanks to the increased diffusion coefficient of the extracellular compartment.

To test this hypothesis, we extended our Monte Carlo simulations to include *T*_2_ relaxations, and set the *T*_2_ values of 18 ms and 36 ms for the intra- and extracellular compartments, respectively (similar to the values measured in yeast by Eriksson et al.). We repeated the simulation with an increased extracellular *T*_2_ value of 100 ms. All parameters are otherwise the same as in Sec. (3 C). We then analyzed the simulated data using the bi-compartmental model, oblivious to the difference in the *T*_2_ values, and calculated the error in the extracted exchange rate.

For the FEXSY experiment, we extracted *τ*_*i*_ = 461 ms, 16% error. Increasing the extracellular *T*_2_ to 100 ms, we got *τ* = 398 ms, 28% error. In contrast, the corresponding exchange rates extracted from the log-linear fit of the CG-PGSTE experiment for the case of *g* = 100 G/cm were 504 ms and 500 ms, a 9% error.

Recall that in the absence of a difference in the *T*_2_ relaxations, the values extracted from the corresponding simulated data were off only by 3% from the in silico exchange rate. We conclude that the difference in *T*_2_ values can, at least partially, explain the quantitative discrepancy between the two methods. Thus, until a more rigorous follow-up is performed, we encourage pairing of exchange and *T*_2_ measurements.

## 6. DISCUSSION

### A. General observations

As water exchange across membranes is both important and difficult to quantify, the main objectives of the present study were to evaluate the ability of diffusion NMR-based methods to do just that in systems of increasing complexity and to compare between available methods. Here, we used two of the more prevalent non-invsasive diffusion NMR-based methods for measuring exchange, namely the CG-PGSTE and FEXSY methods, to study and analyze, using simple modeling approaches, apparent water exchange in yeast cell samples and in the more complex optic nerves both before and after fixation.

The two approaches confirmed previous expectations and demonstrated that when blindly using current methodologies, the apparent intracellular water exchange rates are faster in yeast cells as compared to optic nerves affording intracellular mean residence times (*τ*_*i*_) in the ranges of ∼300-600 and ∼800-4000 ms, respectively. We found the apparent averaged *τ*_*i*_ values extracted from the FEXSY experiments to be smaller than that extracted from the log-linear fit of the CG-PGSTE NMR experiments both for yeast cells and optic nerves. Checking the experimental precision of these methods, the extracted values were found to be quite constant for about 12-14 hours, and good reproducibility was found between samples. However, we found that the extracted values for the optic nerves, in contrast to the yeast cells’ values, appear to strongly depend on the NMR parameters used. In yeast cells, with sufficient diffusion weighting, the extracted intracellular MRT reached stable and constant values. In con-trast, for optic nerves, this occurred only at very high diffusion weighting and to a much lesser degree, rendering the reported values problematic – a meaningful comparison can be made only when the experimental parameters are exactly the same. Furthermore, we found and discussed few indications that the WM signal cannot be analyzed using a bi-compartmental model. This motivated us to attempt fitting the data using a novel tri-compartmental model. Indeed, the new model fit the data significantly better and reports dramatically different exchange rate, with (*τ*_*i*_) in the ranges of ∼400-1000 ms – larger but much closer to the values extracted from yeast cells.

### B. Comparison with literature

#### i. Yeast cells

The exchange rates obtained for yeast cell samples by the two methods used were found to be robust, reproducible, and nearly independent of the parameters used when sufficient diffusion weighting was achieved and when obeying the SGP approximation. Under these conditions, it appears that the approximations used, especially for the log-linear fit of the CG-PGSTE NMR experiments, describe the yeast cells system well. This is also manifested in Figure 3. However, quantitatively, it appears that the average intracellular MRT obtained from the log-linear fit of the CG-PGSTE experiments, 574 ± 6 ms, is greater than the average value extracted from the FEXSY experiments, which was found to be 368 ± 14 ms for the global fit. In several cases, diffusion NMR-based methods were used to study exchange in red blood cells and in neuronal cells [33, 34, 36–38]. However, there are not too many diffusion NMR-based studies of exchange in yeast cells in the literature [32, 46–49, 65, 66]. Our CG-PGSTE results are similar to the values obtained for yeast cells by Tanner assuming two Gaussian compartments [46]. However, intracellular MRTs in our study were found to be somewhat shorter than the average value recently reported by the Baldwin group (∼ 860 ms) assuming that one of the populations is restricted to spheres [65]. Note that therein the effect of *T*_1_ relaxation was also taken into account.

The values we obtained from the FEXSY experiments for *τ*_*i*_ and *τ*, using the global fit were 368 ± 14 ms and 251 ± 2 ms, respectively. These values are similar to the values obtained by Lasič et al. [49] using the same method and using the phenomenological fit. These values are, however, slightly larger than those reported by Eriksson et al. [47] but are slightly smaller than the value reported by the same group in the work of Å slund et al. [46]. When comparing our data with the literature, we restricted ourselves to the data obtained for yeast cells using diffusion NMR-based methods [32, 46– 49, 65, 66, 93, 94]. Note, however, that in different studies different experimental parameters were used, e.g., diffusion weighting, diffusion times, TEs, etc. These differences may affect to some extent the exchange rates and may be responsible for some of the differences in the extracted values. It should be noted that the yeast strain may also have contributed to the variation in the reported values, as demonstrated re-cently [94]. In this recent study, a *k*_*i*_ value of 3.11 s^−1^ was reported from the FEXSY experiment, which is very similar to the value we found for the same yeast strain.

Comparing the FEXSY and CG-PGSTE NMR methods, it appears that the apparent exchange rates extracted from the FEXSY experiments are larger than the exchange rates extracted from the log-linear fit of the CG-PGSTE NMR experiments. Although the origin of this observation is not yet clear, it is important to note that in the only study in which the two methods were used to measure exchange in the same system, i.e., in leukemia K562 cells, the same qualitative trend was observed [72]. However, therein, 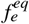 was not extracted from the available FEXSY data, and the comparison was between *τ* of FEXSY and *τ*_*i*_ of CG, so no conclusion can be decisively drawn. There, much faster exchange rates were documented in accordance with previous studies in which water exchange across the membrane in other human cells were measured [33, 34, 36–38].

Interestingly, we found that 15 minutes of fixation of yeast cells significantly affected their exchange rate: The intracellular MRT, as obtained from the log-linear fit of the PGSTE NMR data, decreased from 574 ± 6 to 337 ± 10 ms after fixation. The respective *τ*_*i*_ values extracted from the FEXSY experiments also decreased from 368 ± 14 to 146 ± 24 ms after fixation implying an increase in membrane permeability due to fixation. These results seem to suggest that fixation, generally used to preserve the microstructural features of the sample, significantly affects membrane permeability.

#### ii. Optic nerves – inadequate modeling

The extracted exchange rates and apparent MRTs for the nerve samples were found to be less robust and more dependent on the parameters used in the measurement. Markedly, the CG-PGSTE fit was shown to be highly dependent on the gradient strength, in direct contradiction with the bicompartmental model predictions. The log-linear fit of the CG-PGSTE experiment showed that, as expected, the apparent intracellular MRT (*τ*_*i*_) is larger in nerves as compared to yeast cells. Intracellular MRT values of 2260 ± 69 and 2490 ± 223 ms were extracted from the log-linear fit of the CG-PGSTE experiment for optic nerves before and after fixation, respectively. Note that these values, which are larger than the ones extracted for yeast cells, are also much higher than the values extracted for neuronal cells by the Leibfritz group using a similar methodology [37, 38]. It is important to note, however, as shown in Figure 5 and Tables III and S1 that the validity indices of the approximations used in the loglinear fit of the CG-PGSTE NMR in the case of optic nerves are significantly lower than those observed in the case of yeast cell samples. Nevertheless, the above values are in the vicinity of the numbers extracted more recently in WM using FEXSY and FEXI [49, 51].

Indeed, the values we extracted for the apparent intracellular MRT *τ*_*i*_ from the FEXSY experiment performed on optic nerves before and after fixation were found to be 834 ± 135 ms and 617 ± 235 ms, respectively. As stated above, these values are not very different from the values recently reported by different groups for different WM areas using FEXSY and FEXI [49, 51]. It is important to note, however, that in our hands, the extracted values were found to depend strongly on the parameters used to collect the FEXSY data (e.g., *t*_*m*_, *g*_*max*_, TE). We observe that *τ*_*i*_ values are higher, despite the fact that the extracted *τ* values are relatively small. This occurs since the *τ*_*i*_ values are obtained by dividing *τ* values by 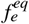 (see Eq. (19)). However, forcing the bi-compartmental fit leads to an inaccurate measurement of 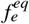.

In addition, Figures 7(b) and S4(b) show that, in contrast to what was found for yeast cells, even at very long mixing times the normalized ADCs do not approach the asymptotic value of 1 as expected but rather plateau around 0.3 − 0.4. We demonstrated that forcing the fit to 1 results in a *τ* which is off by an order of magnitude from the actual value. Leaving a free parameter *c* as in Eq. (35) solves this issue.

As the evidence supporting breakdown of the bicompartmental model in WM accumulated, we decided to consult Monte Carlo simulations.

### C. Simulations

This breakdown of the bi-compartmental model in the case of the optic nerve may be due to the existence of an additional component or components which are filtered out by the “filter block”, but do not recover, since they are simply nonexchanging.

An alternative explanation would be the much smaller difference between *ADC*_*e*_ and *ADC*_*i*_ than that observed in the case of yeast cells, for example. This implies that the ability of the filter block to practically nullify one of the populations while leaving the other nearly intact is much more limited. However, in Figure 9 we present the results of simulations of an HCP lattice, the most dense packing possible, where the extracellular domain is even more restricted than the intracellular domain. Hence, both domains exhibit highly nonGaussian diffusion. Yet, despite the apparent low efficiency of the filter block, the simulated FEXSY experiment shows a return of the normalized ADCs to unity. We can thus rule out this explanation.

In experiments, we observed a discrepancy between the absolute values measured by the methods, a factor of about ∼ 1.5. The simulations of both methods performed equally well, with a very accurate quantitative estimate of the exchange rates. As mentioned, in the only previous experimental comparison, *τ*_*i*_ from both methods were not compared, but rather the *τ* of FEXSY and the *τ*_*i*_ of CG [72]. Here, we compared results of idealized simulations, and demonstrated that both methods allow for very accurate inference of the exchange rates. Markedly, in our simulations we used a compartment size and permeability that are roughly those encountered in yeast cells, and the same q-values, diffusion times and mixing times as in the experiments we performed on the real yeast cells. If these parameters were to cause the discrepancy, we would have observed it in the simulations as well. Since we did not, we can rule out these as possible factors.

Different non-diffusive signal relaxations remain a plausible factor, and in Sec. 5 we showed that at least part of the effect can be attributed to different values of spin-spin relaxations for the two spin populations. A major obstacle in this discussion is the lack of ground truth, which prevents us from determining which method gives a more accurate estimate of the exchange rate.

### D. The lack of ground truth

One of the main problems in developing and challenging new methods and modeling approaches to study water exchange across membranes, especially in biological tissues, is the lack of a ground truth or a measurable system in which the absolute exchange rate is known a priori. Such ground truth is much easier to obtain in the case of microstructural studies, for example [15–17, 73–80].

One way to overcome this difficulty is to measure a specific system, having a specific exchange rate, using different methods and compare the results. Indeed, an effort in that direction was recently reported by Obata et al. [24]. There, the discrepancy between the measured values was difficult to reconcile. This is not surprising since very different methods for measuring exchange may measure somewhat different exchange processes and hence different rate should be expected. This is one of the reasons for our decision, in this study, to use two diffusion NMR-based methods to measure exchange. Interestingly, we found that even with two of the most prevalent diffusion NMR-based methods for measuring exchange we observe some discrepancy in the extracted absolute values. At this point, it is difficult to pin-point the origin of these differences and determine whether or not they are solely due to *T*_2_ differences. Note, however, that the relative values and observed trends are similar for the exchange rates extracted from the two methods. All of these results imply that, in many cases, diffusion NMR-based methods are only suitable for providing apparent exchange rates (perhaps conducting additional D – *T*_2_ correlation measurements can solve this issue). Therefore, it is important to conduct more comparative studies in which the exchange rate is measured by different NMR methods. Such an approach will enable one to better evaluate the significance of the extracted values and determine the limitations of the different approaches used to measure exchange rates, especially in complex systems like optic nerves and white matter tissues.

Since in most cases only AXRs are obtained, it is advisable to use similar methodologies and even standardized experimental parameters to allow for a more direct and meaningful quantitative comparison between the extracted values. Alternatively, it is advisable to work with an adequate model that is robust to a change in the experimental parameters, such as the new model for WM presented herein.

### E. A new model for white matter analysis

Combining the results from porcine optic nerve ex vivo and simulations, we conclude that the most likely explanation for the breakdown of the bi-compartmental model is the existence of a third compartment that is restricted (hence with a nonGaussian diffusion attenuation) and non-exchanging. In Sec. 4 we derive a proper tri-compartmental model and use it to re-analyze the data. The results are promising in that the fit is significantly better, and there are no longer obvious contradictions between model predictions and results.

Importantly, correct fitting resulted in exchange rates that are considerably faster than those that were calculated using the bi-compartmental model. CG-PFG experiments were found to be 730 ± 40 ms and 803 ± 16 ms for optic nerves before and after fixation, respectively. The respective *τ*_*i*_ values extracted from the FEXSY experiments before and after fixation were found to be 530 ± 125 ms and 387 ± 104 ms, respectively. This finding has implications for previous studies.

As mentioned in Sec. 1, Stanisz et al. [40] proposed a tricompartmental model of WM: axons (prolate ellipsoid), glial cells (spheres), and extracellular medium. However, in their model, both restricted compartments were assumed to be exchanging with an unrestricted extracellular medium. Interestingly, a later variation of this model replaced the spheres with a highly restricted and non-exchanging “dot compartment” [95]. Although it is tempting to map between the dot compartment and the third non-exchanging compartment discussed herein, one should note that previous studies estimated its signal fraction to be small (upper limit of 2.7%) [95–97]. Furthermore, Dhital et al. used isotropic diffusion encoding to investigate neuronal tissues and concluded that any substantial third population of WM that is isotropic must have a very small contribution to the signal, if not negligible. In other words, if a third compartment exists, it is likely to be anisotropic (where there might be a certain combination of compartment shape anisotropy and ensemble anisotropy [98]). Note that, because all exchange experiments were performed with a short TE of 9 ms, myelin and bound water may contribute to the third non-exchanging component.

In a comprehensive set of *in silico* experiments, Brusini et al. introduced a WM model with explicit account of exchange through myelin sheets [99]. The exchange rate decreased significantly with larger axonal diameters and a higher number of myelin wraps. Previous experiments have shown that the optic nerve has a large population of small-diameter axons [11], albeit the distribution is broad, and large-diameter axons have a greater contribution to the signal fraction. It is thus plausible that the third compartment is the coarse-grained fraction of large-diameter axons with a high number of myelin wraps, where exchange is completely negligible. Then, the observed exchange takes place between small-diameter axons with a low number of myelin wraps and the extracellular compartment. To further explore this hypothesis, a follow-up experiment could compare between normal and pathological WM. In MS and ALS, for example, the axons are unmyelinated/damaged, or with abnormal axon sizes [6, 7]). If further experiments confirm the hypothesis, one can think of a gen-eralization to the AxCaliber model of Assaf et al. [11]: A bi-compartmental model, with exchange between the extracellular compartment and a heterogeneous cylindrical intracellular compartment, where the axon diameters are drawn from a Gamma distribution. Furthermore, the exchange rate itself is a function of the diameter of the axon.

### F. The effect of fixation

In this work, we quantified the effect of fixation on the transmembrane water exchange rate. We found that fixation generally increases the exchange rate (significantly in yeast and less so in optic nerves). The increase in permeability is consistent with a recent DEXSY study by Williamson et al. [59], and with a relaxation-based study by Shepherd et al. [100]. This can be explained by the formation of methylene cross-links involving membrane proteins, in a way that disrupts the structure of the membrane [77, 100].

Importantly, previous NMR studies on fixation have primarily focused on WM and other neuronal tissues [59, 77, 100–102]. Here, before turning to the optic nerve, we first considered the fixation of yeast cells, which exhibit much simpler microstructural features and are well described by the bi-compartmental model. Furthermore, we used immersion fixation while most previous studies of the CNS performed perfusion fixation [59, 100–102]. This allowed us to disentangle between the effects of formaldehyde on cell culture and those effects that can be possibly attributed to perfusion itself, such as the reduction of tissue volume. In fact, both in yeast and in WM, we observed a larger extracellular fraction after fixation. In yeast, the FEXSY bi-exponential fit (Sec. 3 A ii) reveals an increase of the extracellular fractional population in equilibrium 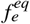 from 0.68 ± 0.03 to 0.79 ± 0.03. This result is in agreement with a recent fluorescence microscopy study, showing that chemical fixation caused on average a 9% decrease in yeast cell diameter [103].

Many previous studies were performed under the assumption that exchange does not take place during the diffusion time, and hence exchange was not incorporated into the modeling of fixation effects [77, 101, 102]. Although this assumption is not unreasonable, in Appendix B we show that improperly neglecting to incorporate exchange can lead to an underestimation of the intracellular fraction and, at the same time, to an overestimation of the compartment size. This warrants caution in future experiments, as fixation is likely a confounding variable that may affect both the exchange rate and the fractional water populations [104].

Lastly, with the tri-compartmental fit, we note that immersion fixation had the same effect of increasing the extracellu-lar fraction from 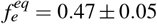 to 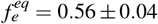. We stress that in our tri-compartmental model 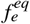 and 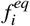 still sum up to 1, and their relative size compared to the third compartment is undetermined, unless we explicitly incorporate the extent of the third compartment attenuation by the filter block. In other words, the reported increase in the extracellular fraction upon fixation is only relative to the intracellular fraction.

### G. Limitations of the used models

The results obtained in the present study clearly demonstrate that the bi-compartmental model describes the yeast cells in an acceptable manner. Similarly, a tri-compartmental model seems to better fit WM data. In the simple modeling approaches used here, the effect of relaxation was ignored altogether. In addition, no information of the geometry, the anisotropy [105–107], size distribution, and orientation dispersion, which are all much more important and significant in the optic nerve, were included in the modeling and analysis. Importantly, Khateri et al. rely on simulations to demonstrate that there may be another source of exchange that should be accounted for: diffusion-mediated exchange, where water molecules diffuse between anisotropic domains with different orientations relative to the measurement gradient direction [107]. A similar idea can be found in a recent tricompartmental model of gray matter, with diffusion-mediated exchange between dendritic shaft and spines [108].

### H. Recent developments to notice

Very recently, using extremely high static field gradients and state-of-the-art analysis and processing techniques to measure extremely fast exchange rates, Williamson et al. showed promising preliminary results that set diffusion-NMR as a candidate to directly measure cellular metabolic activity via the rate constant for water exchange across cell membranes (without exogenous tracers or contrast agents) [59]. Going beyond basic research, the immediate obstacle appears to be the high field gradients in use, which exceed clinically available gradients by orders of magnitude. This study also explores systematically active vs. passive mechanisms for transmembrane water exchange, an important step forward. In this context, it is further demonstrated on an oxygen–glucose deprivation model of stroke that exchange measurements can provide functional information [59] that is additional to the classic apparent diffusion coefficient (ADC) measurement [109]. The last point is supported by another very recent work, using a different d-MR method, diffusion time–dependent diffusion kurtosis imaging, DKI(*t*). Therein, Wu et al. demonstrate that hypoxic-ischemic brain injury increases transmembrane water exchange [110].

The application of exchange measurements using d-MRS is not limited to transmembrane water exchange. A new study quantifies the exchange in the liquid-liquid phase separation (LLPS) phenomenon, namely the exchange dynamics between the condensed and dispersed phases [67]. While the theoretical considerations and experimental techniques are equivalent to those used in measuring transmembrane exchange rate, new studies [67, 111] further position d-MRS as a non-invasive tool for measuring certain chemical reaction rates (where the two reacting species or states differ sufficiently in their mobilities).

As an example, Zhou et al. used d-MRS to measure dynamic ligand exchange on colloidal nanocrystals, namely the adsorption-desorption rates of ligands binding to colloids [111]. Markedly, they used a probe with a maximal gradient strength of 3000 G/cm. Recall that high gradients were also employed in Williamson’s study, reaching 1530 G/cm. With higher gradients, faster exchange rates become accessible for d-MRS measurement [59, 71, 111]. Here we have shown that, at least for yeast cells where the bi-compartmental model holds, changing the gradient strength in the range 15 − 250 G/cm does not have a significant effect on the extracted exchange rate. Hence, it is safe to say that the same exchange process is measured. It would be interesting to conduct an experiment in which the diffusion weighting is varied over a larger range, to verify that different processes can indeed be measured, and to identify them.

Since MRI is non-invasive and has a high penetration depth, an ongoing effort is underway to develop efficient and nontoxic MRI gene reporters that can track gene expression in vivo. Most techniques rely on certain contrast agents that affect signal relaxation through spin-spin or lattice-spin interactions [112, 113], or alternatively, use contrast agents for Chemical Exchange Saturation Transfer (CEST) Imaging [114, 115]. In 2016, Mukherjee et al. established that human aquaporines, and specifically AQP1, can be utilized as diffusion-MRI reporter genes [116], paving the way for a potential new family of gene reporters. Recently, they have systematically reassured safety concerns about AQP1 overexpression in diverse cell types in vitro [117]. In their work, the increased membrane permeability to water molecules due to AQP1 overexpression was measured by a simple diffusion weighted imaging technique. As the contrast is explicitly due to transmembrane water exchange, we believe that it could be worthwhile to apply specialized diffusion-MRI techniques, such as those presented here, for the direct quantification of exchange and permeability in such experiments. Recall that FEXI has already been employed in this context by Schilling et al. They have used the urea transporter as a gene reporter, relying on transporter mediated increase in plasma membrane water exchange [53].

## 7. CONCLUSIONS

We used two of the most prevalent diffusion NMR-based methods for studying apparent exchange rates in yeast cells and optic nerves both before and after fixation. The data was analyzed based on the simple log-linear fit of the CG-PGSTE experiment and the more recently developed FEXSY method. We found that simple modeling of the data obtained from the above MR diffusion-based methods enable extracting the intracellular MRT in the yeast cell samples. The log-linear fit of the PGSTE experiments yielded *τ*_*i*_ values of about 554 ± 6 ms and 337 ± 10 ms for yeast cells before and after fixation, respectively. Interestingly, with the FEXSY experiments we extracted *τ*_*i*_ values of 368 ± 14 ms and 146 ± 24 ms, respectively, for yeast cells before and after fixation. The origin of the discrepancy between the absolute values remains unclear, but we show that it can be attributed, at least partially, to different spin-spin relaxation in the intra- and extracellular compartments.

For optic nerves, the log-linear fit of the PGSTE experiment yielded *τ*_*i*_ values of about 2260 ± 69 ms and 2490 ± 223 ms before and after fixation. With FEXSY *τ*_*i*_ values of 834 ± 135 ms and 617 ± 235 ms before and after fixation, respectively, demonstrating that fixation increases the exchange rates also in optic nerves but to a much lesser degree.

We also presented indications that the bi-compartmental model is inadequate for WM, and used Monte Carlo simulation to further support this hypothesis. It appears that if one insist on using such models, only apparent values can be extracted in complex systems such as optic nerves and other WM tissues, and these are very sensitive the experimental parameters used.

Based on analysis of the experimental data and simulations, we showed that in WM tissues like the optic nerve the signal can be better described by a model which is at least tricompartmental, with the additional compartments not partaking in exchange. We have discussed possible candidates for such a compartment and the prospect of more elaborate modeling that accounts for the axonal-diameter distribution. Furthermore, the compartment should be small enough so that the spins diffusing in it are restricted. We derived the proper model and corroborated it with data and simulations. With this model for optic nerves, the CG-PGSTE experiment yielded *τ*_*i*_ = 730 ± 40 ms, and FEXSY yielded *τ*_*i*_ = 530 125 ms. These values are considerably smaller than those extracted via what seems to be an erroneous bi-compartmental fit of WM, and are in fact much closer to values measured for the yeast sample (where the latter is indeed bi-compartmental).

Lastly, it is extremely important to perform more comparative studies in which the exchange rate is measured in the same system using different methods and perhaps analyzed with different models. Moreover, we argue that constructing an agreed upon gold standard with a known exchange rate to corroborate experimental results is of outmost importance.

## Abbreviations used

ADC: apparent diffusion coefficient
ALS: amyotrophic lateral sclerosis
AXR: apparent exchange rate
CG: constant gradient
CNS: central nervous system
DEXSY: diffusion exchange spectroscopy
FEXSY: filter-exchange spectroscopy
FEXI: filter-exchange imaging
HCP: hexagonal close-packed
MS: multiple sclerosis
MR: magnetic resonance
MRS: magnetic resonance spectroscopy
MRI: magnetic resonance imaging
MRT: mean residence time
NMR: nuclear magnetic resonance
PFG: pulse-field gradient
PGSE: pulsed gradient spin echo
PGSTE: pulsed gradient stimulated echo
RF: radiofrequency
SGP: short gradient pulse
SNR: signal to noise ratio
WMR: white matter

## Acknowledgments

YC wishes to thank the Israel Science Foundation (ISF, Jerusalem, Israel) for a grant that supported, partially, the purchase of the 9.4 T MR instrument used in this project. YC wishes to thank the U.S.-Israel Binational Science Foundation (BSF), grant number 2019-25. This project has received funding from the European Research Council (ERC) under the European Union’s Horizon 2020 research and innovation program (grant agreement No. 947731 to S.R.).

## Data availability statement

The data that supports the findings of this study are available in the supplementary material of this article.

## Appendix A

**FIG. S1.**
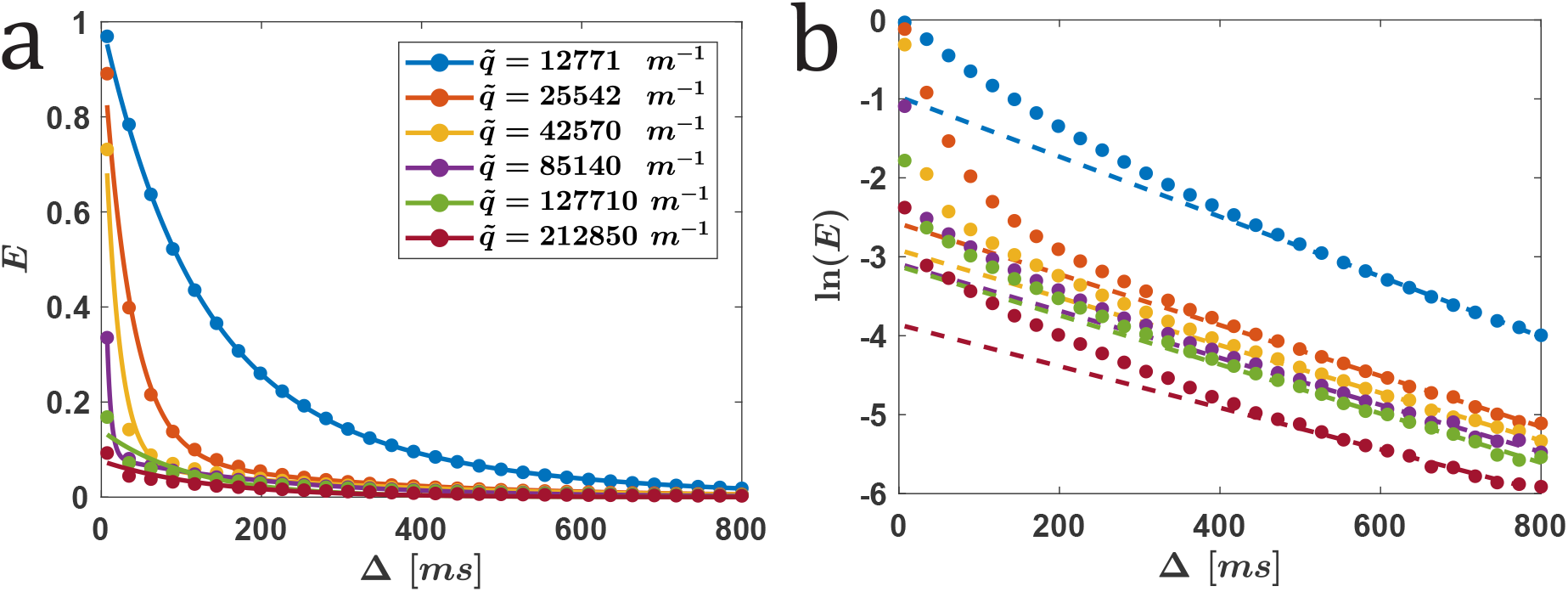
CG-PGSTE NMR experiments on fixed yeast cells: (a) Normalized signal *E* vs. the diffusion time (full circles). Data was fitted with Eq. (6) (lines), and (b) ln(*E*) vs. the diffusion time (full circles). Data was fitted with Eq. (16) using the ten data points with the highest diffusion times (dashed lines). Data is shown for different q-values obtained with *δ*_*eff*_ = 2 ms and g-values of 15 (blue), 30, 50, 100, 150 and 250 G/cm.

**FIG. S2.**
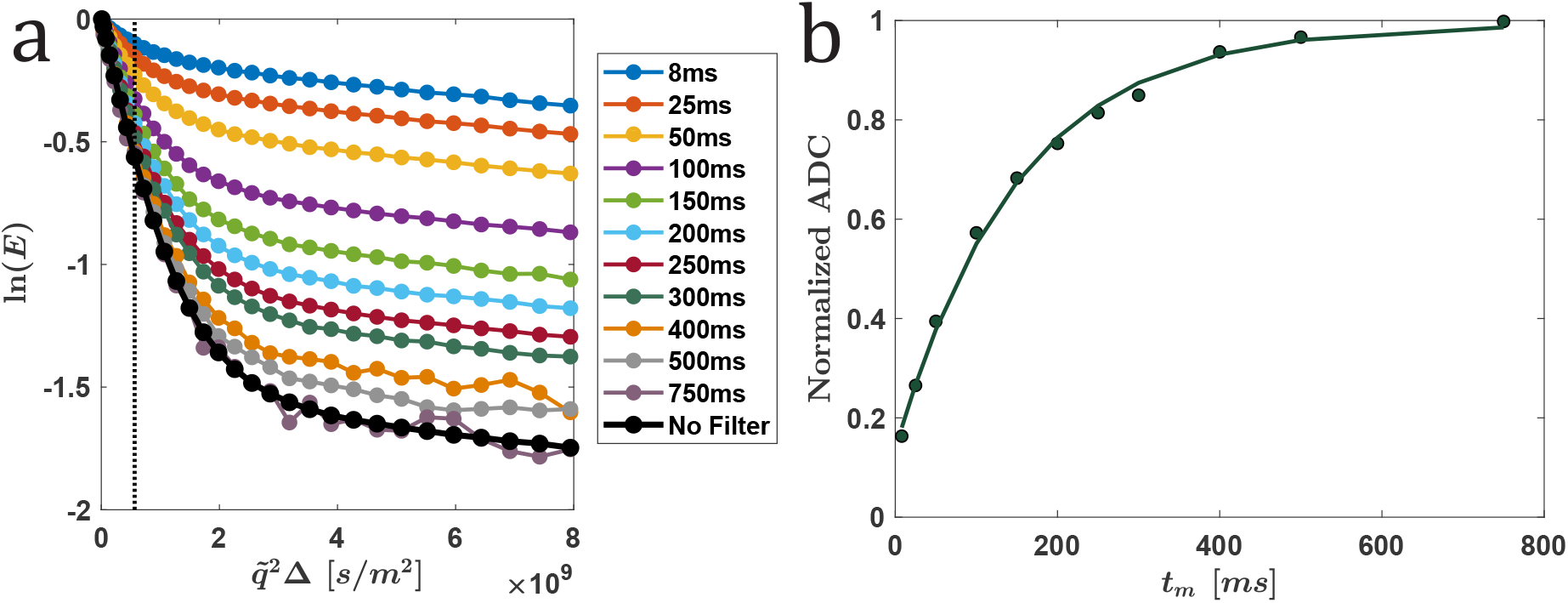
Results of a FEXSY experiment performed on a fixed yeast cell sample: (a) The attenuation of the natural logarithm of the normalized echo ln(*E*) vs. 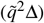 for different *t*_*m*_ values of the FEXSY and the no-filter experiment (full circles), and (b) normalized ADCs extracted (full circles) from the data presented in (a). Note that the gradient strength of the filter was 150 G/cm and that the gradients in the measurement block were incremented from 2.5 to 150 G/cm in 30 steps. The effective *δ* was 2 ms. Normalized ADCs were computed from ADC extracted from the first 8 points of each graph in (a) divided by the ADC extracted from the no filter experiment (ADC_*eq*_). The lines in (a) are just to guide the eyes while the line in (b) is a fit to Eq. (23).

**FIG. S3.**
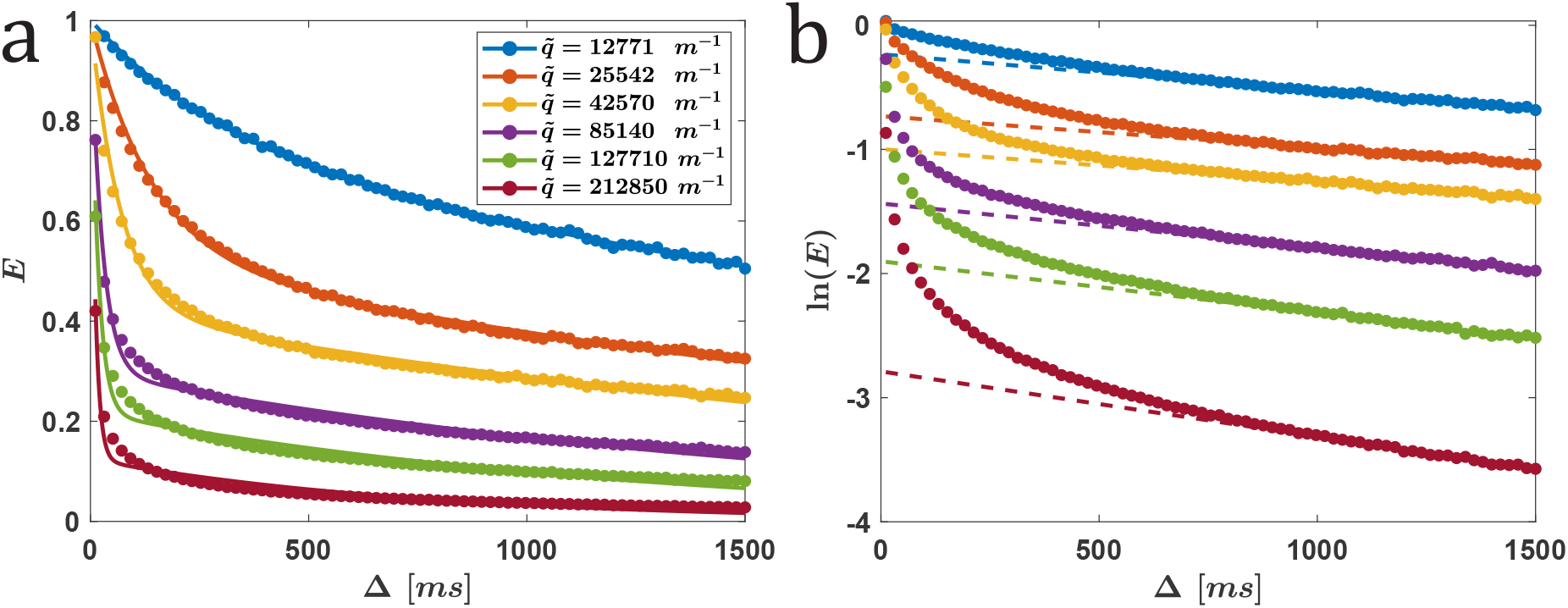
CG-PGSTE NMR experiments on a fixed optic nerve: (a) Normalized signal (*E*) vs. the diffusion time (full circles). Data was fitted with Eq. (6) (lines), and (b) ln(*E*) vs. the diffusion time (full circles). Data was fitted with Eq. (16) using the 20 data points (out of 76 collected) with the highest diffusion times (dashed lines). Data is shown for different q-values obtained with *δ*_*eff*_ = 2 ms and g-values of 15 (blue), 30, 50, 100, 150 and 250 G/cm.

**FIG. S4.**
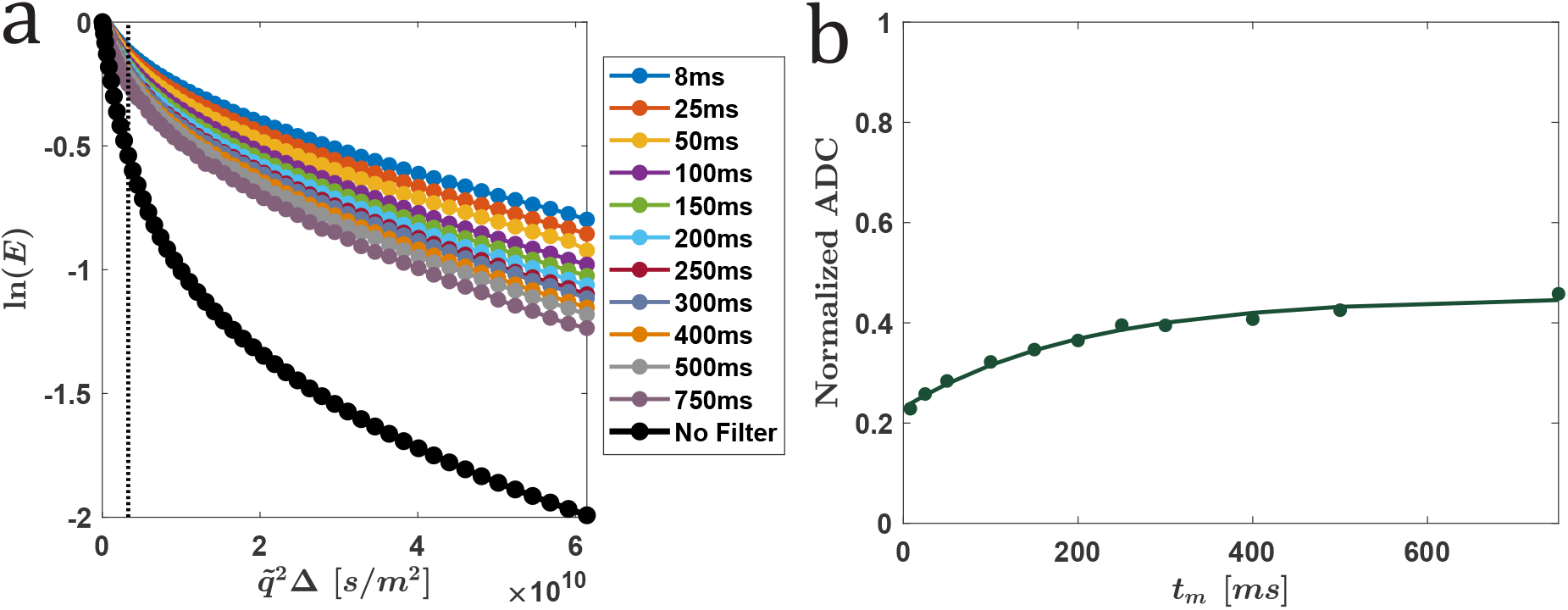
Results of a FEXSY experiment performed on a fixed optic nerve sample: (a) The natural logarithm of the normalized echo ln(*E*) vs. 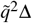 for different *t*_*m*_ values of the FEXSY and the non-filter experiments (full circles), and (b) normalized ADCs extracted (filled circle) from the data presented in (a). Note that the gradient strength of the filter was 250 G/cm and that the gradients in the measurement block were incremented from 2.5 to 250 G/cm in 52 steps. The length of the sine-shaped gradient pulses was 3.14 ms, i.e., *δ*_*eff*_ = 2 ms. For both filter and measurement PGSTE blocks Δ/*δ*_*eff*_ of 35/2 ms were used and *t*_*m*_ was varied between 8 and 750 ms. Normalized ADCs were computed from ADCs extracted from the first 12 points of each graph in (a) divided by the ADC extracted from the no filter experiment (ADC_*eq*_). The lines in (a) are just to guide the eyes while the line in (b), is a fit to the phenomenological model (Eq. (23)).

**TABLE S1.** An exemplary porcine optic nerve sample. Fitted *τ*_*i*_ values (ms) and *R*^2^ for different gradient strengths and effective diffusion times *δ*_eff_. The letter ‘a’ stands for data that could not be fitted because of the poor signal quality (low SNR) at these high q-values.

| Gradient strength<br>(G/cm) | $\delta_{eff} = 4$ ms | | $\delta_{eff} = 6$ ms | | $\delta_{eff} = 10$ ms | |
| --- | --- | --- | --- | --- | --- | --- |
| | $\tau_i$ (ms) | $R^2$ | $\tau_i$ (ms) | $R^2$ | $\tau_i$ (ms) | $R^2$ |
| 15 | 3788 | 0.9627 | 2873 | 0.9807 | 2462 | 0.9897 |
| 30 | 2840 | 0.9772 | 2234 | 0.9784 | 2218 | 0.9595 |
| 50 | 2226 | 0.9718 | 2135 | 0.9754 | 2095 | 0.9382 |
| 100 | 1871 | 0.9698 | 1738 | 0.9297 | 1878 | 0.7540 |
| 150 | 1734 | 0.9641 | 1731 | 0.7806 | a | 0.5> |
| 250 | 1895 | 0.9356 | a | 0.5> | a | 0.5> |

**FIG. S5.**
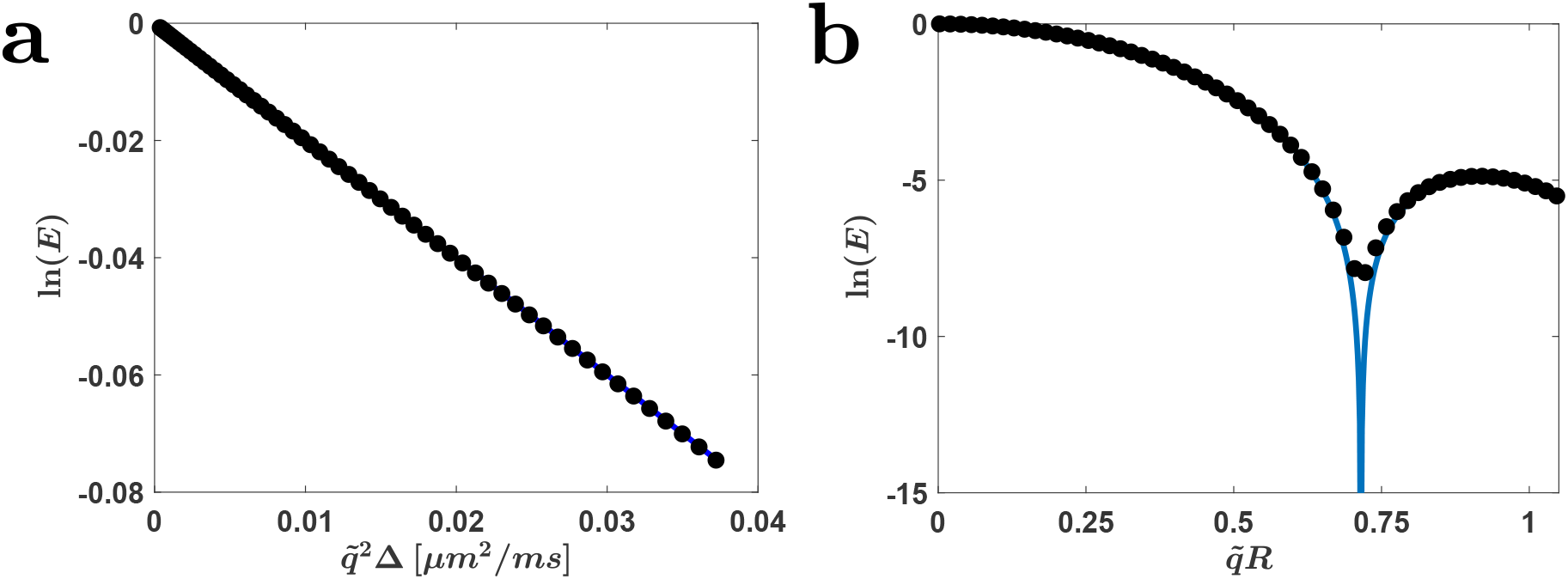
Simulating the two limiting cases of *κ* → ∞ and *κ* → 0. Full circles come from Monte-Carlo simulations while the solid lines are the theoretical predictions. (a) *κ* → ∞: A free diffusion attenuation curve, which is Gaussian according to Eq. (5). We set *D* = 2 *µ*m^2^*/*ms and simulated 10^5^ spins with the simulation time step d*t* = 1 *µ*s. Here we plot ln(*E*) vs. 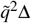 such that the slope gives the diffusion coefficient. Linear fit results in *D* = 2.003 *µ*m^2^*/*ms. (b) *κ* → 0: Diffusion restricted to a sphere. We simulated 10^7^ spins with d*t* = 1 *µ*s. We set Δ = 100 ms, *R* = 5 *µ*m and *D* = 2 *µ*m^2^*/*ms, such that *ξ* ≫ 1 and we anticipate a diffraction pattern according to Eq. (9).

**TABLE S2.** Results for the CG-PFG simulation in Sec. 4 C. The fit was performed according to Eq. (34). For each gradient strength, Δ was varied linearly up to 800 ms with a thousand points. For the analysis points 400-750 were taken, since these points produced overall best *R*^2^s. The highest gradient strength consistently shows a poor fit.

| Gradient Strength<br>(G/cm) | Tri-compartmental fit |  |  |  |
| --- | --- | --- | --- | --- |
| | $\tau_i$ (ms) | A | B | $R^2$ |
| 15 | 548 | 0.06 | 0.78 | 0.9995 |
| 30 | 573 | 0.18 | 0.37 | 0.9996 |
| 50 | 628 | 0.25 | 0.08 | 0.9994 |
| 100 | 536 | 0.17 | 0.09 | 0.9983 |
| 150 | 513 | 0.10 | 0.09 | 0.9950 |
| 250 | 228 | 0.02 | 0.09 | 0.7634* |

**FIG. S6.**
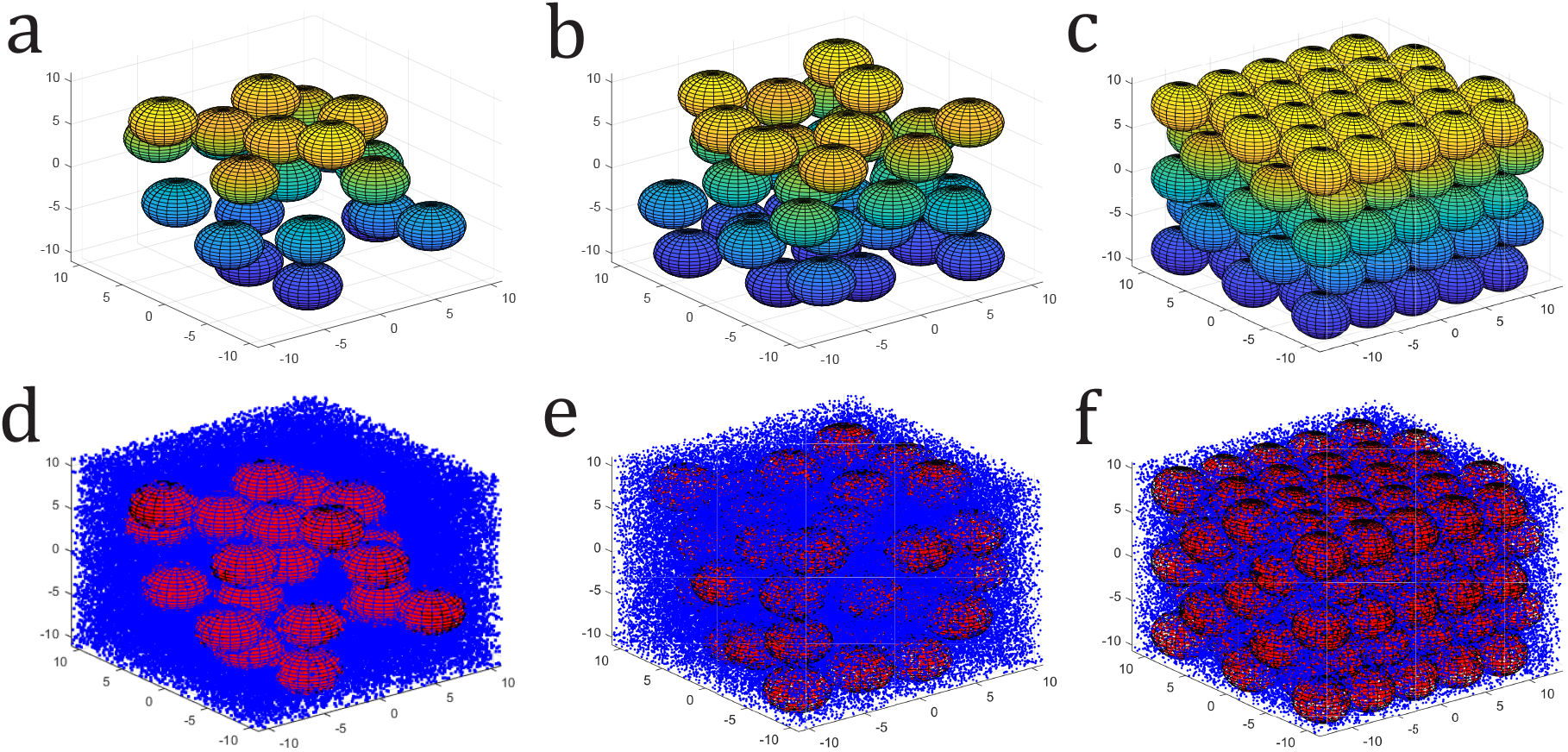
Monte-Carlo simulations of water molecules (spins) propagating in a domain containing permeable spherical cells (with radius of 5 *µ*m). Top row, shows the positions of the distributed spheres for the cases of (a) 23 randomly distributed spheres, (b) 43 randomly distributed spheres and (c) hexagonal close packing (HCP). The bottom row, panels (d)-(f), are the corresponding plots of the initial positions of the spins inside the domains. Spins that are in the intracellular domain are colored red, whilst spins that are in the extracellular domain are colored blue.

**FIG. S7.**
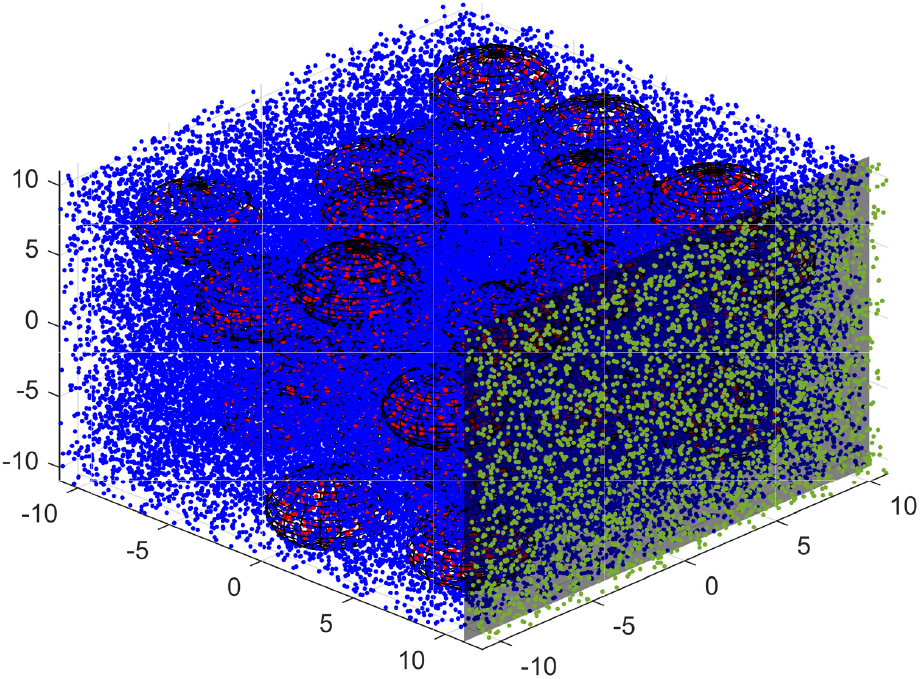
The same domain as in Figure S6, with an additional non-exchanging compartment. The width of the extension in the measurement axis was set to 2 *µ*m to ensure restriction. The additional spins in the third compartment are colored green.

## Appendix B

**How excluding exchange affects model accuracy?**

To assess if and to what extent exclusion of exchange from the bi-compartmental model leads to extraction of erroneous values, we conducted a set of simulations of simple PFG experiments, where we vary the exchange rate. We repeated the procedure for different values of diffusion time Δ. We then analyzed the results using a simple model of two non-exchanging populations, according to Eq. (10), where 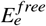 is given by Eq. (5) and 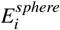 is given by Eq. (9), and we recall that *f* = 1 − *f*. The extracted *R* and *f*_*e*_ values are summarized in Tables S3 and S4 and discussed at the end of Sec. 3 C.

**TABLE S3.**
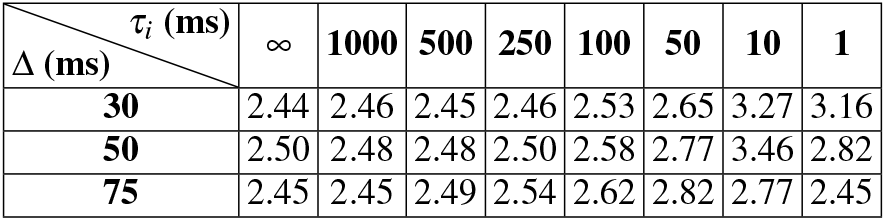
The extracted cell radius in *µ*m as function of Δ and *τ*_*i*_. We set *R* = 2.5 *µ*m, *D* = 2 *µ*m^2^*/*ms and simulated 10^6^ spins with the simulation time step d*t* = 1 *µ*s.

| $\Delta$ (ms) \ $\tau_i$ (ms) | ∞ | 1000 | 500 | 250 | 100 | 50 | 10 | 1 |
| --- | --- | --- | --- | --- | --- | --- | --- | --- |
| 30 | 2.44 | 2.46 | 2.45 | 2.46 | 2.53 | 2.65 | 3.27 | 3.16 |
| 50 | 2.50 | 2.48 | 2.48 | 2.50 | 2.58 | 2.77 | 3.46 | 2.82 |
| 75 | 2.45 | 2.45 | 2.49 | 2.54 | 2.62 | 2.82 | 2.77 | 2.45 |

**TABLE S4.** The extracted *f*_*e*_ as function of Δ and *τ*_*i*_. We set *R* = 2.5 *µ*m, *D* = 2 *µ*m^2^*/*ms and simulated 10^6^ spins with the simulation time step d*t* = 1 *µ*s. The actual extracellular fractional population in the simulation was *f*_*e*_ = 0.74.

| $\Delta$ (ms) \ $\tau_i$ (ms) | ∞ | 1000 | 500 | 250 | 100 | 50 | 10 | 1 |
| --- | --- | --- | --- | --- | --- | --- | --- | --- |
| 30 | 0.70 | 0.71 | 0.72 | 0.73 | 0.75 | 0.79 | 0.90 | 0.94 |
| 50 | 0.70 | 0.72 | 0.73 | 0.75 | 0.80 | 0.84 | 0.93 | 0.96 |
| 75 | 0.72 | 0.73 | 0.74 | 0.77 | 0.83 | 0.89 | 0.84 | 0.98 |

An interesting observation: A model that does not account for exchange vastly underestimates the number of restricted spins that contribute to this signal. It can lead to an estimate of *f*_*i*_ that is orders of magnitude smaller than its real value, giving the wrong impression that the intracellular compartment is almost devoid of spins.

## Notes

### Competing Interest Statement

The authors have declared no competing interest.

### Summary of Updates

Revised version. More Discussion. More references. Proofing. Slight title change.

## References

[1] W.S. Price, NMR studies of translational motion, Cambridge University Press, Cambridge, 2009.

[2] P.T. Callaghan, Translational dynamics and magnetic resonance, Oxford University Press, Oxford, 2011.

[3] D.K. Jones (Ed.), Diffusion MRI: Theory, methods, and applications, Oxford University Press, Oxford, 2010.

[4] J. Kärger, D.M. Ruthven, D.N. Theodorou, Diffusion in nanoporous materials, Wiley-VCH-Verlag GmbH & Co. KGaA, 2012.

[5] J.M. Ritchie (1982). On the relation between fibre diameter and conduction velocity in myelinated nerve fibres. Proc. Roy. Soc. London Ser. B: Biol.Sci. 217:29–35.

[6] S. Cluskey, D.B. Ramsden (2001). Mechanisms of neurodegeneration in amyotrophic lateral sclerosis. Mol. Pathol. 54:386–392.

[7] N. Evangelou, D. Konz, M.M. Esiri, S. Smith, J. Palace, P.M. Matthews (2002). Size-selective neuronal changes in the anterior optic pathways suggest a differential susceptibility to injury in multiple sclerosis. J. Neuro-Ophthalmology. 22:143.

[8] Y. Assaf, A. Mayk, Y. Cohen (2000). Displacement imaging of spinal cord using q-space diffusion-weighted MRI. Magn. Reson. Med. 44:713–722.

[9] Y. Cohen, Y. Assaf (2002). High b-value q-space analyzed diffusion-weighted MRS and MRI in neuronal tissues - A technical review. NMR Biomed. 15:516–542.

[10] Y. Assaf, P.J. Basser (2005). Composite hindered and restricted model of diffusion (CHARMED) MR imaging of the human brain. Neuroimage. 27:48–58.

[11] Y. Assaf, T. Blumenfeld-Katzir, Y. Yovel, P.J. Basser (2008). AxCaliber: A method for measuring axon diameter distribution from diffusion MRI. Magn. Reson. Med. 59:1347–1354.

[12] D.C. Alexander (2008). A general framework for experiment design in diffusion MRI and its application in measuring direct tissue-microstructure features. Magn. Reson. Med. 60:439–448.

[13] H. Zhang, T. Schneider, C.A. Wheeler-Kingshott, D.C. Alexander (2012). NODDI: Practical in vivo neurite orientation dispersion and density imaging of the human brain. Neuroimage. 61:1000–1016.

[14] X. Wang, M.F. Cusick, Y. Wang, P. Sun, J.E. Libbey, K. Trinkaus, R.S. Fujinami, S.K. Song (2014). Diffusion basis spectrum imaging detects and distinguishes coexisting subclinical inflammation, demyelination and axonal injury in experimental autoimmune encephalomyelitis mice. NMR Biomed. 27:843–852.

[15] Y. Cohen, D. Anaby, D. Morozov (2017). Diffusion MRI of the spinal cord: from structural studies to pathology. NMR Biomed. 30(3):e3592.

[16] D.C. Alexander, T.B. Dyrby, M. Nilsson, H. Zhang (2017). Imaging brain microstructure with diffusion MRI: practicality and applications. NMR Biomed. 32(4):e3841.

[17] D.S. Novikov, E. Fieremans, S.N. Jespersen, V.G. Kieselev (2018). Quantifying brain microstructure with diffusion MRI: Theory and parameter estimation. NMR Biomed. 32(4):e3998.

[18] J. Veraart, D. Nunes, U. Rudrapatna, E. Fieremans, D.K. Jones, D.S. Novikov, N. Shemesh (2020). Noninvasive quantification of axon radii using diffusion MRI. elife. 9:e49855.

[19] S.J. Chen, J.F. Yang, F.P. Kong, J.L. Ren, K. Hao, M. Li, Y. Yuan, X.C. Chen, R.S. Yu, J.F. Li, G. Leng, X.Q. Chen, J.Z. Du (2014). Over-activation of corticotropin releasing factor receptor type 1 and aquporin-4 by hypoxia indices cerebral edema. Proc. Natl. Acad. Sci. USA. 111:13199–13204.

[20] Z. He, X. Wang, Y. Wu, J. Jia, Y. Hu, X. Yang, J. Li, M. Fan, L. Zhang, J. Guo, M.C.P. Leung (2014). Treadmill pretraining ameliorate brain edema in ischemic stroke via down–regulation of aquaporin-4: An MRI study in rats. PLoS ONE. 9:e84602.

[21] J.O. Breen-Norris, B. Siow, C. Walsh, B. Hipwell, I. Hill, T. Roberts, M.G Hall, M.F Lythgoe, A. Ianus, D.C. Alexander, S. Walker-Samuel (2020). Measuring diffusion exchange across the cell membrane with DEXSY (Diffusion Exchange Spectroscopy). Magn. Reson. Med. 84:1543–1551.

[22] D. Shi, S. Li, F. Liu, X. Jiang, L. Wu, L. Chen, Q. Zheng, H. Bao, H. Guo, J. Xu (2024). Comprehensive characterization of tumor therapeutic response with simultaneous mapping cell size, density, and transcytolemmal water exchange. arXiv:2408.01918.

[23] D. Shi, F. Liu, S. Li, L. Chen, X. Jiang, J.C. Gore, Q. Zheng, H. Guo, J. Xu (2024). Restriction-induced time-dependent transcytolemmal water exchange: Revisiting the Kärger exchange model. J. Magn. Reson. 367:107760.

[24] T. Obata, J. Kershaw, T. Tachibana, T. Miyauchi, Y. Abe, S. Shibata, H. Takuwa, I. Aoki, M. Yasui (2018). Comparison of diffusion-weighted MRI and anti-Stokes Raman scattering (CARS) measurements of the inter-compartmental exchange-time of water in expression-controlled aquaporin-4 cells. Sci. Rep. 8:17954.

[25] D.S. Grebenkov (2014). Exploring diffusion across permeable barriers at high gradients. II. Localization regime. J. Magn. Reson. 248:164–176.

[26] J. Kärger (1969). Zur bestimmung der diffusion in einem zweibereich-system mit hilfe von gespulsten feldgaradienten. Ann. Der Phys. 24:1–4.

[27] J. Kärger (1985). NMR self-diffusion studies in heterogeneous systems. Adv. Colloid Interface Sci. 23:129–148.

[28] J. Kärger, H. Pfeifer, W. Heink (1988). Principles and application of self-diffusion measurements by nuclear magnetic resonance. Adv. Magn. Reson. 12:1–89.

[29] D. Wijesekera, T. Stait-Gardner, A. Gupta, J. Chen, G. Zheng, A.M. Torres, W.S. Price (2019). A complete derivation of the Kärger equations for analysing NMR diffusion measurements of exchanging systems, Concepts Magn. Reson. Part A. 47A:c21468

[30] E.O. Stejskal, J.E. Tanner (1965). Spin diffusion measurements: Spin echoes in the presence of a time-dependent field gradient. J. Chem. Phys. 42:288–292.

[31] J.E. Tanner (1970). Use of the stimulated echo in NMR diffusion studies. J. Chem. Phys. 52:2523–2526.

[32] J.E. Tanner (1978). Transient diffusion in a system partitioned by permeable barriers- application to NMR measurements with a pulsed field gradient. J. Chem. Phys. 69:1748–1754.

[33] J. Andrasko (1976). Water diffusion permeability of human erythrocytes studied by pulsed gradient NMR techniques. Bioch. Biophys. Acta. 428:304–311.

[34] A.R. Waldeck, P.W. Kuchel, A.J. Lennon, B.E. Chapman (1997). NMR diffusion measurements to characterize membrane transport and solute binding, Prog. NMR Spectrosc. 30:39–68.

[35] W.S. Price, A.V. Barzykin, K. Hayamizu, M. Tachiya. A model for diffusive transport through a spherical interface probed by pulsed-field gradient NMR (1998). Biophys. J. 74:2259–2271.

[36] J. Pfeuffer, U. Flögel, D. Leibfritz (1998). Monitoring of cell volume and water exchange time in perfused cells by diffusion weighted 1H NMR spectroscopy. NMR Biomed. 11:11–18.

[37] C. Meier, W. Dreher, D. Leibfritz (2003). Diffusion in compartmental systems: I. A comparison of an analytical model with simulations., Magn. Reson. Med. 50:500–509.

[38] C. Meier, W. Dreher, D. Leibfritz (2003). Diffusion in compartmental systems. II. diffusion-weighted measurements of rat brain tissue in vivo and postmortem at very large b-values. Magn. Reson. Med. 50:510–514.

[39] A. Szafer, J. Zhong, J.C. Gore (1995). Theoretical model for water diffusion in tissues. Magn. Reson. Med. 33:697–712.

[40] G.J. Stanisz, A. Szafer, G.A. Wright, R.M. Henkelman (1997). An analytical model of restricted diffusion in bovine optic nerve. Magn. Reson. Med. 37:103–111.

[41] P.T. Callaghan, M.E. Komlosh (2002). Locally anisotropic motion in a macroscopically isotropic system: displacement correlations measured using double pulsed gradient spin-echo NMR. Magn. Reson. Chem. 40:S15–S19.

[42] P.T. Callaghan, S. Godefroy, B. Ryland (2003). Use of the second dimension in PGSE NMR studies of porous media. Magn. Reson. Imaging. 21:243–248.

[43] P.T. Callaghan, I. Furó (2004). Diffusion-diffusion correlation and exchange as a signature for local order and dynamics. J. Chem. Phys. 120:4032–4038.

[44] T.X. Cai (2023). In search of lost time: development of rapid magnetic resonance methods to probe time-varying diffusion (Doctoral dissertation, University of Oxford).

[45] D. Benjamini, M.E. Komlosh, P.J. Basser (2017). Imaging local diffusive dynamics using diffusion exchange spectroscopy MRI. Phys. Rev. Letters. 118:158003.

[46] I. Åslund, A. Nowacka, M. Nilsson, D. Topgaard (2009). Filter exchange PGSE NMR determination of cell membrane permeability. J. Magn. Reson. 200:291–295.

[47] S. Eriksson, K. Elbing, O. Söderman, K. Lindkvist-Petersson, D. Topgaard, S. Lasič (2017). NMR quantification of diffusional exchange in cell suspensions with relaxation rate differences between intra- and extracellular compartments. PloS One. 12:e0177273.

[48] Y. Scher, S. Reuveni, Y. Cohen (2020). Constant-gradient FEXSY: An efficient method for measuring exchange. J. Magn. Reson. 311:106667.

[49] S. Lasič, M. Nilsson, J. Lätt, F. Ståhlberg, D. Topgaard (2011). Apparent exchange rate mapping with diffusion MRI. Magn. Reson. Med. 66:356–365.

[50] M. Nilsson, J. Lätt, D. Van Westen, S. Brockstedt, S. Lasič, F. Sttåhlberg, D. Topgaard (2013). Noninvasive mapping of water diffusional exchange in the human brain using filter-exchange imaging. Magn. Reson. Med. 69:1573–1581.

[51] S. Lasič, S. Oredsson, S.C. Partridge, L.H. Saal, D. Topgaard, M. Nilsson, K. Bryskhe (2016). Apparent exchange rate for breast cancer characterization, NMR Biomed. 29:631–639.

[52] B. Lampinen, F. Szczepankiewicz, D. van Westen, E. Englund, P.C. Sundgren, J. Lätt, F. Ståhlberg, M. Nilsson (2017). Optimal experimental design for filter exchange imaging: Apparent exchange rate measurement in the healthy brain and in intracranial tumors, Magn. Reson. Med. 77:1104–1114.

[53] F. Schilling, S. Ros, D.E. Hu, P. D’Santos, S. McGuire, R. Mair, A.J. Wright, E. Mannion, R.J.M. Franklin, A.A. Neves, K.M. Brindle (2017). MRI measurements of reporter-mediated increases in transmembrane water exchange enable detection of a gene reporter. Nat. Biotech. 35:75–80.

[54] S. Lasič, A. Chakwizira, H. Lundell, C.F. Westin, M. Nilsson (2024). Tuned exchange imaging: Can the filter exchange imaging pulse sequence be adapted for applications with thin slices and restricted diffusion?. NMR Biomed. 37:e5208.

[55] D. Benjamini, P.J. Basser (2016). Use of marginal distributions constrained optimization (MADCO) for accelerated 2D MRI relaxometry and diffusometry. J. Magn. Reson. 271:40–45.

[56] T.X. Cai, D. Benjamini, M.E. Komlosh, P.J. Basser, N.H. Williamson (2018). Rapid detection of the presence of diffusion exchange. J. Magn. Reson. 297:17–22.

[57] N.H. Williamson, R. Ravin, T.X. Cai, D. Benjamini, M. Falgairolle, M.J. O’Donovan, P.J. Basser (2020). Real-time measurement of diffusion exchange rate in biological tissue. J. Magn. Reson. 317:106782.

[58] T.X. Cai, N.H. Williamson, R. Ravin, P.J. Basser (2022). Disentangling the effects of restriction and exchange with diffusion exchange spectroscopy. Front. Phys. 10:805793.

[59] N.H. Williamson, R. Ravin, T.X. Cai, M. Falgairolle, M.J. O’Donovan, P.J. Basser (2023). Water exchange rates measure active transport and homeostasis in neural tissue. PNAS Nexus. 2:pgad056.

[60] T.X. Cai, N.H. Williamson, R. Ravin, P.J. Basser (2024). The Diffusion Exchange Ratio (DEXR): A minimal sampling of diffusion exchange spectroscopy to probe exchange, restriction, and time-dependence. J. Magn. Reson. 366:107745.

[61] R. Song, Y.Q. Song, M. Vembusubramanian, J.L. Paulsen (2016). The robust identification of exchange from T2-T2 time-domain features, J. Magn. Reson. 265:164–171.

[62] A. Ordinola, S. Cai, P. Lundberg, R. Bai,, E. Özarslan (2023). On the sampling strategies and models for measuring diffusion exchange with a double diffusion encoding sequence. Magn. Reson. Lett. 3:232–247.

[63] I.O. Jelescu, A. de Skowronski, F. Geffroy, M. Palombo, D.S. Novikov (2022). Neurite exchange imaging (NEXI): A minimal model of diffusion in gray matter with inter-compartment water exchange, NeuroImage. 256:119277.

[64] J.L. Olesen, L. Østergaard, N. Shemesh, S.N. Jespersen (2022). Diffusion time dependence, power-law scaling, and exchange in gray matter. NeuroImage. 251:118976.

[65] G. Karunanithy, R.J. Wheeler, L.R. Tear, N.J. Farrer, S. Faulkner, A.J. Baldwin (2019). INDIANA: An in-cell diffusion method to characterize the size, abundance and permeability of cells. J. Magn. Reson. 302:1–13.

[66] M. Schillmaier, A. Kaika, G.J. Topping, R. Braren, F. Schilling (2023). Repeatability and reproducibility of apparent exchange rate measurements in yeast cell phantoms using filter-exchange imaging. Magn. Reson. Mater. Phy. 36(6):957–974.

[67] M. Novakovic, N. Han, N.C. Kathe, Y. Ni, L. Emmanouilidis, F.H.T. Allain (2025). LLPS REDIFINE allows the biophysical characterization of multicomponent condensates without tags or labels. Nat. Commun. 16:4628.

[68] C. Li, E. Fieremans, D.S. Novikov, Y. Ge, J. Zhang (2023). Measuring water exchange on a preclinical MRI system using filter exchange and diffusion time dependent kurtosis imaging. Magn. Reson. Med. 89:1441–1455.

[69] X. Jiang, S.P. Devan, J. Xie, J.C. Gore, J. Xu (2022). Improving MR cell size imaging by inclusion of transcytolemmal water exchange. NMR Biomed. 35:e4799.

[70] N. Shemesh, S.N. Jespersen, D.C. Alexander, Y. Cohen, I. Drobnjak, T.B. Dyrby, J. Finsterbusch, M.A. Koch, T. Kuder, F. Laun, M. Lawrenz, H. Lundell, P.P. Mitra, M. Nilsson, E. Özarslan, D. Topgaard, C.F. Westin (2016). Conventions and nomenclature for double diffusion encoding NMR and MRI. Magn. Reson. Med. 75:82–87.

[71] D.S. Grebenkov, D. Van Nguyen, J.R. Li (2014). Exploring diffusion across permeable barriers at high gradients. I. Narrow pulse approximation. J. Magn. Reson. 248:153–163.

[72] X. Tian, H. Li, X. Jiang, J. Xie, J.C. Gore, J. Xu (2017). Evaluation and comparison of diffusion NMR methods for measuring apparent transcytolemmal water exchange rate constant. J. Magn. Reson. 275:29–37.

[73] L. Avram, Y. Assaf, Y. Cohen (2004). The effect of rotational angle and experimental parameters on the diffraction patterns and micro-structural information obtained from q-space diffusion NMR: Implication for diffusion in white matter fibers. J. Magn. Reson. 169:30–38.

[74] E. Fieremans, Y. De Deene, S. Delputte, M.S. Özdemir, Y. D’Asseler, J. Vlassenbroeck, K. Deblaere, E. Achten, I. Lemahieu (2008). Simulation and experimental verification of the diffusion in an anisotropic fiber phantom. J. Magn. Reson. 190:189–199.

[75] B. Siow, I. Drobnjak, A. Chatterjee, M.F. Lythgoe, D.C. Alexander (2012). Estimation of pore size in a microstructure phantom using the optimised gradient waveform diffusion weighted NMR sequence. J. Magn. Reson. 214:51–60.

[76] D. Morozov, L. Bar, N. Sochen, Y. Cohen (2013). Modeling of the diffusion MR signal in calibrated model systems and nerves. NMR Biomed. 26:1787–1795.

[77] S. Richardson, B. Siow, E. Panagiotaki, T. Schneider, M.F. Lythgoe, D.C. Alexander (2014). Viable and fixed white matter: Diffusion magnetic resonance comparisons and contrasts at physiological temperature. Magn. Reson. Med. 72:1151–1161.

[78] J. Xu, H. Li, K.D. Harkins, X. Jiang, J. Xie, H. Kang, M.D. Does, J.C. Gore (2014). Mapping mean axon diameter and axonal volume fraction by MRI using temporal diffusion spectroscopy. Neuroimage. 103:10–19.

[79] D. Morozov, L. Bar, N. Sochen, Y. Cohen (2015). Microstructural information from angular double-pulsed-field-gradient NMR: From model systems to nerves. Magn. Reson. Med. 74:25–32.

[80] S. Vellmer, D. Edelhoff, D. Suter, I.I. Maximov (2017). Anisotropic diffusion phantoms based on microcapillaries. J. Magn. Reson. 279:1–10.

[81] E. Fieremans, H.H. Lee (2018). Physical and numerical phantoms for the validation of brain microstructural MRI: A cookbook, Neuroimage. 181:39–61.

[82] J. Cohen-Adad (2018). Microstructural imaging in the spinal cord and validation strategies. Neuroimage. 182:169–183.

[83] H. Li, X. Jiang, J. Xie, J.C. Gore, J. Xu (2017). Impact of transcytolemmal water exchange on estimates of tissue microstructural properties derived from diffusion MRI. Magn. Reson. Med. 77:2239–2249.

[84] P.T. Callaghan (1995). Pulsed-gradient spin-echo NMR for planar, cylindrical and spherical pores under conditions of wall relaxation. J. Magn. Reson. Ser. A. 113:53–59.

[85] P.T. Callaghan, Principles of nuclear magnetic resonance microscopy. Clarendon press, 1993.

[86] I. Åslund, D. Topgaard (2009). Determination of the selfdiffusion coefficient of intracellular water using PGSE NMR with variable gradient pulse length. Magn. Reson. Med. 201:250–254.

[87] B. Balinov, B. Jonsson, P. Linse, O. Soderman (1993). The NMR self-diffusion method applied to restricted diffusion. Simulation of echo attenuation from molecules in spheres and between planes. J. Magn. Reson. Ser. A. 104:17–25.

[88] N. Shemesh, E. Özarslan, P.J. Basser, Y. Cohen (2012). Accurate noninvasive measurement of cell size and compartment shape anisotropy in yeast cells using double-pulsed field gradient MR. NMR Biomed. 25(2):236–246.

[89] E. Fieremans, D.S. Novikov, J.H. Jensen, J.A. Helpern (2010). Monte Carlo study of a two-compartment exchange model of diffusion. NMR Biomed. 23:711–724.

[90] J.E. Tanner (1983). Intracellular diffusion of water. Arch. Biochem. Biophys. 224:416–428.

[91] S.N. Jespersen, M. Pedersen, H. Stødkilde-Jørgensen (2005). The influence of a cellular size distribution on NMR diffusion measurements. Eur. Biophys. J. 34:890–898.

[92] S.Y. Huang, T. Witzel, B. Keil, A. Scholz, M. Davids, P. Dietz, E. Rummert, R. Ramb, J.E. Kirsch, A. Yendiki, Q. Fan, Q. Tian, G. Ramos-Llordén, H. Lee, A. Nummenmaa, B. Bilgic, K. Setsompop, F. Wang, A.V. Avram, M. Komlosh, D. Benjamini, K.N. Magdoom, S. Pathak, W. Schneider, D.S. Novikov, E. Fieremans, S. Tounekti, C. Mekkaoui, J. Augustinack, D. Berger, A. Shapson-Coe, J. Lichtman, P.J. Basser, L.L. Wald, B.R. Rosen (2021). Connectome 2.0: Developing the next-generation ultra-high gradient strength human MRI scanner for bridging studies of the micro-, meso-and macroconnectome. NeuroImage. 243:118530.

[93] N. Moutal, M. Nilsson, D. Topgaard, D.S. Grebenkov (2018). The Kärger vs bi-exponential model: Theoretical insights and experimental validations. J. Magn. Reson. 296:72–78.

[94] M. Soltesova, H. Elicharova, P. Srb, M. Ruzicka, L. Janisova, H. Shychrova, J. Lang (2019). Nuclear magnetic resonance investigation of water transport through the plasma membrane of various yeast species. FEMS Microb. Lett. 399:fnz220.

[95] C.M. Tax, F. Szczepankiewicz, M. Nilsson, D.K. Jones (2020). The dot-compartment revealed? Diffusion MRI with ultra-strong gradients and spherical tensor encoding in the living human brain. Neuroimage. 210:116534.

[96] B. Dhital, E. Kellner, V.G. Kiselev, M. Reisert (2018). The absence of restricted water pool in brain white matter. Neuroimage. 182:398–406.

[97] M. Palombo, A. Ianus, M. Guerreri, D. Nunes, D.C. Alexander, N. Shemesh, H. Zhang, (2020). SANDI: A compartment-based model for non-invasive apparent soma and neurite imaging by diffusion MRI. Neuroimage. 215:116835.

[98] N. Shemesh, Y. Cohen (2011). Microscopic and compartment shape anisotropies in gray and white matter revealed by angular bipolar double-PFG MR. Magn. Reson. Med. 65(5):1216–1227.

[99] L. Brusini, G. Menegaz, M. Nilsson (2019). Monte Carlo simulations of water exchange through myelin wraps: implications for diffusion MRI. IEEE Trans. Med. Imag. 38(6):1438–1445.

[100] T.M. Shepherd, P.E. Thelwall, G.J. Stanisz, S.J. Blackband (2009). Aldehyde fixative solutions alter the water relaxation and diffusion properties of nervous tissue. Magn. Reson. Med. 62(1):26–34.

[101] C. Wang, L. Song, R. Zhang, F. Gao (2018). Impact of fixation, coil, and number of excitations on diffusion tensor imaging of rat brains at 7.0 T. Eur. Radiol. Exp. 2(1):25.

[102] T. Santini, A. Shim, J.J. Liou, N. Rahman, G. Varela-Mattatall, M.D. Budde, W. Inoue, S. Everling, C.A. Baron (2025). Investigating microstructural changes between in vivo and perfused ex vivo marmoset brains using oscillating gradient and b-tensor encoded diffusion MRI at 9. 4 T. Magn. Reson. Med. 93(2):788–802.

[103] D. Appadurai, L. Gay, A. Moharir, M.J. Lang, M.C. Duncan, O. Schmidt, D. Teis, T.N. Vu, M. Silva, E.M. Jorgensen, M. Babst (2020). Plasma membrane tension regulates eisosome structure and function. Mol. Biol. Cell. 31(4):287–303.

[104] A. Chakwizira, C.F. Westin, J. Brabec, S. Lasič, L. Knutsson, F. Szczepankiewicz, M. Nilsson (2023). Diffusion MRI with pulsed and free gradient waveforms: effects of restricted diffusion and exchange. NMR Biomed. 36(1):e4827.

[105] C. K. Sønderby, H.M. Lundell, L.V. Søgaard, T.B. Dyrby (2014). Apparent exchange rate imaging in anisotropic systems. Magn. Reson. Med. 72(3):756–762.

[106] H.G. Shin, X. Li, H.Y. Heo, L. Knutsson, F. Szczepankiewicz, M. Nilsson, P.C. van Zijl (2024). Compartmental anisotropy of filtered exchange imaging (FEXI) in human white matter: What is happening in FEXI?. Magn. Reson. Med. 92(2):660–675.

[107] M. Khateri, M. Reisert, A. Sierra, J. Tohka, V.G. Kiselev (2022). What does FEXI measure?. NMR Biomed. 35:e4804.

[108] K. Şimşek, A. Chakwizira, M. Nilsson, M. Palombo (2025). The role of dendritic spines in water exchange measurements with diffusion MRI: Time-Dependent Single Diffusion Encoding MRI. arXiv:2506.18229.

[109] M.E. Moseley, Y. Cohen, J. Mintorovitch, L. Chileuitt, H. Shimizu, J. Kucharczyk, M.F. Wendland, P.R. Weinstein (1990). Early detection of regional cerebral ischemia in cats: comparison of diffusion-and T2-weighted MRI and spectroscopy. Magn. Reson. Med. 14:330–346.

[110] D. Wu, H.H. Lee, R. Ba, V. Turnbill, X. Wang, Y. Luo, P. Walczak, E. Fieremans, D.S. Novikov, L.J. Martin, F.J. Northington, J. Zhang (2024). In vivo mapping of cellular resolution neuropathology in brain ischemia with diffusion MRI. Sci. Adv. 10:eadk1817.

[111] X. Zhou, Z. Pang, W. Cao, Z. Cao, J. Zhu, Y. Qi, J. Zhu, X. Kong (2022). Diffusion NMR for measuring dynamic ligand exchange on colloidal nanocrystals. Anal. Chem. 95:792–801.

[112] A.Y. Louie, M.M. Hüber, E.T. Ahrens, U. Rothbächer, R. Moats, R.E. Jacobs, S.E. Fraser, T.J. Meade. (2000). In vivo visualization of gene expression using magnetic resonance imaging. Nat. Biotechnol. 18(3):321–325.

[113] B. Cohen, K. Ziv, V. Plaks, T. Israely, V. Kalchenko, A. Harmelin, L.E. Benjamin, M. Neeman (2007). MRI detection of transcriptional regulation of gene expression in transgenic mice. Nat. Med. 13(4):498–503.

[114] A. Bar-Shir, G. Liu, Y. Liang, N.N. Yadav, M.T. McMahon, P. Walczak, S. Nimmagadda, M.G. Pomper, K.A. Tallman, M.M. Greenberg, P.C.M. van Zijl, J.W.M. Bulte, A.A. Gilad (2013). Transforming Thymidine into a magnetic resonance imaging probe for monitoring gene expression. J. Am. Chem. Soc. 135(4):1617–1624.

[115] A. Bar-Shir, G. Liu, K.W. Chan, N. Oskolkov, X. Song, N.N. Yadav, P. Walczak, M.T. McMaho, P.C.M. van Zijl, J.W.M. Bulte, A.A. Gilad (2014). Human protamine-1 as an MRI reporter gene based on chemical exchange. ACS Chem. Biol. 9(1):134–138.

[116] A. Mukherjee, D. Wu, H.C. Davis, M.G. Shapiro (2016). Noninvasive imaging using reporter genes altering cellular water permeability. Nat. Commun. 7(1):13891.

[117] A.D.C. Miller, S.P. Chowdhury, H.W. Hanson, S.K. Linderman, H.I. Ghasemi, W.D. Miller, B.M. Gardner, A. Mukherjee (2024). Engineering water exchange is a safe and effective method for magnetic resonance imaging in diverse cell types. J. Biol. Eng. 18(1):30.

